# Epigenomic mapping identifies a super-enhancer repertoire that regulates cell identity in bladder cancers through distinct transcription factor networks

**DOI:** 10.1101/2022.01.11.475197

**Authors:** Hélène Neyret-Kahn, Jacqueline Fontugne, Xiang Yu Meng, Clarice S. Groeneveld, Luc Cabel, Tao Ye, Elodie Guyon, Clémentine Krucker, Florent Dufour, Elodie Chapeaublanc, Audrey Rapinat, Daniel Jeffery, Yann Neuzillet, Thierry Lebret, David Gentien, Irwin Davidson, Yves Allory, Isabelle Bernard-Pierrot, François Radvanyi

## Abstract

Muscle-invasive bladder cancer (BLCA) is an aggressive disease. Consensus BLCA transcriptomic subtypes have been proposed, with two major Luminal and Basal subgroups, presenting distinct molecular and clinical characteristics. However, how these distinct subtypes are regulated remains unclear. We hypothesized that epigenetic activation of distinct super-enhancers could drive the transcriptional programs of BLCA subtypes.

Through integrated RNA-sequencing and epigenomic profiling of histone marks in primary tumours, cancer cell lines, and normal human urothelia, we established the first integrated epigenetic map of BLCA and demonstrated the link between subtype and epigenetic control. We identified the repertoire of activated super-enhancers and highlighted Basal, Luminal and Normal-associated SEs. We revealed the super-enhancer-regulated networks of candidate master transcription factors for Luminal and Basal subgroups including FOXA1 and ZBED2 respectively. FOXA1 CRISPR-Cas9 mutation triggered a shift from Luminal to Basal phenotype, confirming its role in Luminal identity regulation and induced ZBED2 overexpression. In parallel, we showed that both FOXA1 and ZBED2 play concordant roles in preventing inflammatory response in cancer cells through STAT2 inhibition.

Our study furthers the understanding of epigenetic regulation of muscle-invasive BLCA and identifies a co-regulated network of super-enhancers and associated transcription factors providing potential targets for the treatment of this aggressive disease.

## Introduction

Bladder cancer is the tenth most common cancer worldwide, accounting for nearly two thousand cancer-related deaths globally in 2018 (1). Urothelial carcinoma is classified as non-muscle-invasive bladder cancer (NMIBC comprising carcinoma *in situ*, and the pTa and pT1 stages) or the aggressive muscle-invasive bladder cancer (MIBC, stages pT2 to pT4), depending on the level of invasion into the bladder wall (2). Molecular classifications of bladder carcinomas have been established using mainly gene expression profiling studies (3–8). A recent consensus classification of MIBC presents six subtypes, from which tumours can be coarsely divided into two subgroups: the Luminal and the non-Luminal subgroups. Luminal subgroup comprises three Luminal subtypes (LumU, LumNS and LumP) whereas Basal-Squamous subtype (Ba/Sq) constitutes the major part of the non-luminal subgroup (3). Luminal tumours, accounting for about 50% of MIBCs, present high expression of urothelial differentiation markers (GATA3, FOXA1, KRT20, uroplakins) and are enriched in activating mutations of *FGFR3*. Basal tumours also called Basal/Squamous are particularly aggressive and account for ∼35% of MIBCs (3). They are characterized by the overexpression of markers of the basal layer of the urothelium (including KRT5, KRT6), the under-expression of markers of luminal differentiation and activation of EGFR (9). Concerning the NMIBC tumours, the recent UROMOL studies group them into 4 classes including Class 1 associated with luminal differentiation and good prognosis, and a Class 2a comprising high risk tumours (7, 8). One hypothesis to explain the establishment of the different subtypes and their potential plasticity, is that each subtype harbours a regulatory network in which various upstream genomic and epigenomic alterations lead to the activation of a core set of master transcription factors (TFs) that then determine a transcriptomic downstream program. While transcriptional regulators of urothelial differentiation, such as FOXA1, GATA3 and PPARG, have been established as key regulators of the Luminal phenotypes, the essential transcription regulators driving the Ba/Sq subtype have not been elucidated (10–14).

Recent studies have demonstrated that altered enhancer activity drives changes in cell identity and oncogenic transformation, notably through large clusters of highly active enhancers called super-enhancers (SEs) (15–17). Indeed, targeting SE-driven oncogenesis has become a novel therapeutic approach with the advent of BRD4 inhibitors, which inhibit SE activation (18). By regulating the expression of a small number of master TFs, SEs can orchestrate cell- or cancer-specific transcriptional programs. The gold standard for identifying SEs is histone mark profiling (17). The ENCODE roadmap, that profiled histone marks in normal and cancer cell lines, has become a valuable source of information to uncover chromatin organisation, alteration, and subsequent regulation of master regulators but did not include bladder models (19). Recently, two studies provided new insights with the profiling of particular histone marks in bladder cancer samples and cell lines (20, 21). Here, we further characterized bladder cancer epigenetic by integrating transcriptomic and epigenomic profiling of multiple histone marks in human bladder tumours, bladder cancer cell lines, and primary cultures from normal urothelia to produce a comprehensive bladder cancer epigenetic map. With this map, we demonstrated the link between molecular subtype and the underlying epigenetic landscape. Through H3K27ac analysis, we established a repertoire of SEs that are specific to distinct subgroups (Luminal, Ba/Sq subtypes, as well as Normal primary cells), highlighting SE-associated genes with subgroup-specific clinical relevance. From there, we identified the core SE-regulated networks of master TFs that distinguish luminal and basal subgroups, including known and new candidate master TFs. Finally, through functional knock-down and knock-out experiments, we revealed that one of these master TFs (FOXA1) is a key factor in subtype determination antagonized by ZBED2, and that both FOXA1 and ZBED2 present the ability to dampen inflammatory response. Overall, this work provides new data characterizing epigenetic regulation in bladder cancer. We reveal important genes that can be essentials for maintenance of bladder cancer cell identity and present potential new targets to treat aggressive bladder cancers.

## Results

### Integrated bladder cancer chromatin landscape

To elucidate the contribution of chromatin landscape in bladder cancer biology, we generated ChIP-seq for active (H3K27ac) and repressive histone marks (H3K27me3, H3K9me3) in 24 bladder samples (Fig.1). In order to distinguish features of the non-cancerous stromal cells and of normal urothelial cells, we used not only human primary tumours (n=15) from the CIT (*Carte d’Identité des Tumeurs*) cohort (9, 22), but also cellular models (7 bladder cancer cell lines) and patient-derived Normal Human Urothelium in proliferation (NHU, n=2). Of note, tumors were macrodissected to enrich for bladder cancer content (Fig. 1, Fig. S1A). Of the 15 primary tumours, we included 13 MIBCs and 2 NMIBCs to assess the stage-dependence of our results (Fig. 1A, Fig. S1A, and Table S1). With the aim of identifying subtype-specific epigenetic alterations/characteristics, we coupled our ChIP-seq with RNA-seq from the same extraction and classified them according to the current consensus subtypes (3). Of the 13 MIBCs, 2 classified as stroma-rich, 4 classified as basal/squamous (Ba/Sq) and 7 as luminal, including 1 luminal papillary (LumP), 3 luminal unstable (LumU) and 2 luminal non-specified (LumNS). Of the 7 cell lines, 3 were classified Ba/Sq, 3 LumP and one could not be classified (Table S1)(23). Using the recent UROMOL classifier, the two NMIBC samples classified as class 3 (7, 8). Further analysis using subtype deconvolution (WISP, (24)), and previously described regulatory signatures (3, 7), revealed that one of the tumours originally classified as Ba/Sq (T391) was composed of a mixed population of LumP and Ba/Sq cells (Fig. S1B, S1C). Knowing it’s important intra-tumoral heterogeneity, tumour T391 was excluded from differential analyses between subgroups. Peak calling using MACS showed that ChIP-seq for H3K27ac gave the most homogeneous and highest number of peaks across the 24 samples (Fig. S1D).

**Figure 1:**
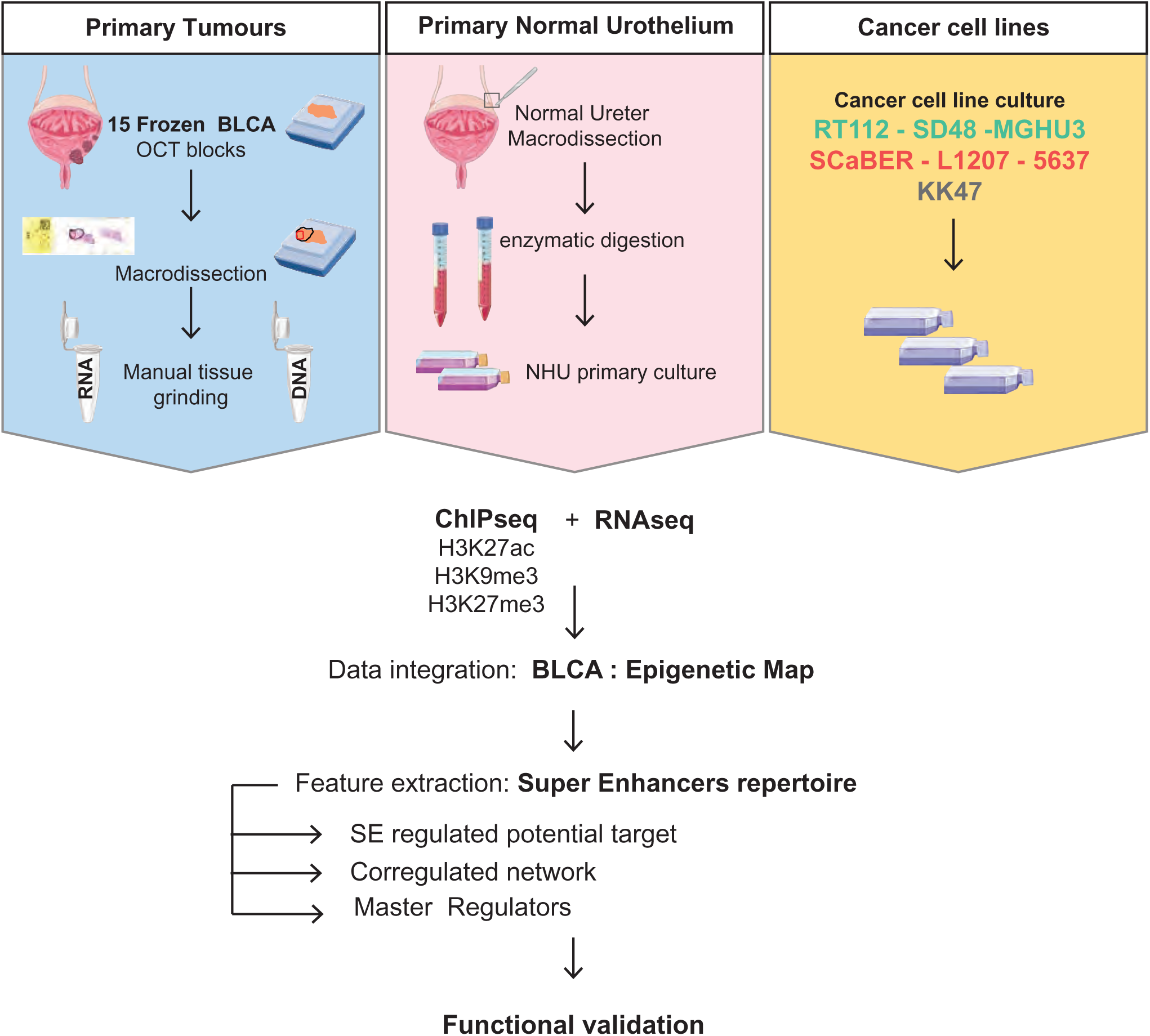
Methodology/Workflow.

We integrated our multi-factorial ChIP-seq profiles using ChromHMM (25), reporting the first integrated epigenetic map in bladder cancer in both primary tumour samples and cell lines (Fig. 2A). Six chromatin states (E1–E6) were assigned according to histone mark enrichments, as previously described (ENCODE, Roadmap project (19)), where H3K27ac-enriched regions correspond to active promoters and enhancers (E2), H3K27me3 and H3K9me3-enriched states associate with repression (E4) or heterochromatin (E6), and regions enriched in both active and repressive marks define bivalent enhancers or promoters (E3). Regions without any marks or only weak H3K9me3 enrichment were designated as quiescent/no marks (E1) or quiescent/weakly repressed (E5), respectively (Fig. 2A, Fig. S2A, B). Analysis of associated RNA-seq data confirmed that gene expression correlates with the expected chromatin states (Fig. 2B). Briefly, genes with transcription start sites (TSSs) in E2 states (active enhancers / promoters) have the highest expression levels, followed by those in E3 states (bivalent enhancers / promoters). Minimal expression was noted for genes with TSSs in the remaining states.

**Figure 2:**
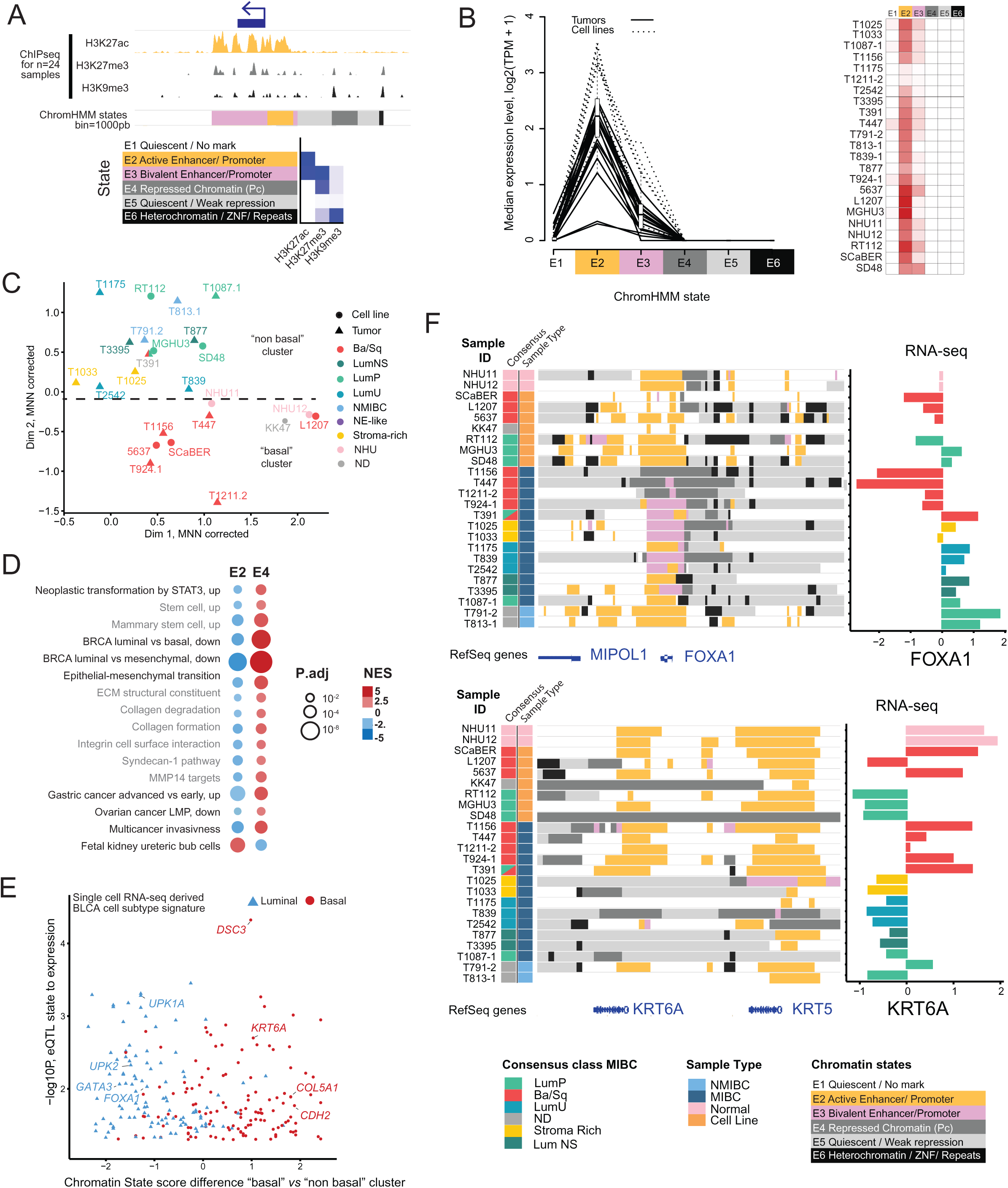
**Chromatin states classify bladder cancers by subgroups** A. ChromHMM principle example and emission order dividing genome in 6 states based on combination of H3K27ac, H3K27me3 and H3K9me3 marks. B. Expression level association with each chromatin state and each sample. C. Two chromatin state clusters revealed by unsupervised analysis of top 1% varying regions using MDS for dimension reduction plus MNN for batch effect correction. D. GSEA functional enrichment analysis of the genes mapped to the MCA Dim2 contributing features. A negative NES indicates significant enrichment in lower Dim2 coordinates (basal direction), and the reverse is in higher Dim2 coordinates (Luminal direction). E. Luminal versus basal tumour cell signature genes identified with single cell RNA-seq analysis showing concordant enrichment in chromatin state clusters. F. Genome Browser view of chromatin states at *FOXA1* and *KRT6* loci with corresponding RNAseq (VST normalized scaled expression).

### Chromatin states classify bladder cancers by transcriptomic subgroups

Next, we sought to classify our samples based on chromatin states for comparison with molecular subtypes. To do this we first performed an unsupervised analysis to select the most distinguishing features from the chromatin profiles (see methods, Fig. S2C) and plotted all samples by multiple correspondence analysis on the most varying regions (MCA, Fig. S2D). Similar to Principal Component Analysis (PCA), but adapted for categorical data, this method of dimensionality reduction separates samples in 2D space by proximity according to the primary (Dim 1) and secondary (Dim 2) dimensions. Thus, greater differences in chromatin profiles are represented by greater distances in the 2D plot. Dim 1 distinguished primary tumour samples from cell lines, which could be indicative of chromatin changes associated with cell culture or stroma content. Interestingly, Dim 2 distinguished Non-basal from Basal subgroups. To confirm this distinction of molecular subtypes, we re-assessed the data with an alternative dimensionality reduction method (MDS), coupled to a batch effect-like correction (MNN), which eliminated most of the cell line vs primary tumour differences while maintaining and strengthening the distinction between Non-Basal and Basal subgroups along Dim 2 (Fig. 2C). Therefore, we identified two clusters derived from differences in chromatin state that are associated with molecular subtypes; a “basal cluster” containing all Ba/Sq samples (except the mixed T391), and a “non-basal cluster” including all luminal, stroma-rich and NMIBC samples (Fig. 2C). Interestingly, NHU cells were located at the border between the two groups (Fig. 2C). To explore the biological pathways associated with the chromatin profiles that could distinguish Luminal from Basal bladder cancers, we ranked genes based on the MCA outputs for Dim 2 and performed Gene Set Enrichment Analysis (GSEA (26, 27), Fig. 2D). As expected, for the basal cluster, we found that active chromatin (E2) was strongly enriched at genes involved in decreased Luminal differentiation, while repressive chromatin (E4) was strongly depleted for these genes (Fig. 2D). Interestingly, genes involved in increased tumour aggressiveness, stemness, extracellular matrix, epithelial-mesenchymal transition and invasion were also enriched in active chromatin and depleted for repressive chromatin in the basal cluster (Fig. 2D). Taking an alternative approach, we derived Basal and Luminal gene signatures from an independent publicly available scRNA-seq dataset (GSM4307111) and compared these genes with the chromatin states associated with the basal and non-basal clusters identified in Fig. 2C (Fig. 2E, see methods). Luminal signature genes were enriched in active state in the non-basal chromatin cluster while Basal signature genes were enriched in active state in the basal chromatin cluster, suggesting epigenetic regulation of signature genes involved in urothelial differentiation. We further illustrate this relationship with two well-described markers of bladder cancer subtypes: *FOXA1* and *KRT6A* (Fig. 2F). FOXA1 expression is higher in Luminal than Basal tumours (11, 14, 28, 29). In agreement, our results showed that *FOXA1* was marked with active (E2) chromatin in Luminals (including LumP, LumNS and LumU), NMIBC samples, and even NHU cells, but harboured repressive chromatin in Ba/Sq tumour samples (Fig. 2F). On the other hand, *KRT6A*, commonly expressed in Basal tumours (3), had active chromatin marks in seven out of eight Ba/Sq samples, as well as NHU, but not in any Luminal sample. Taken together, these results demonstrate the importance of histone marks in the regulation of gene expression driving cell identity in bladder cancer.

### Identification of the bladder super-enhancer repertoire and subtype specificities

To determine if chromatin profiles identify SEs that could control bladder cancer subtype, we determined and annotated typical enhancer and SE regions in our samples using the ROSE algorithm (Table S2), which calls enhancers and particularly super enhancers according to H3K27ac signal (18, 30). Despite signal correction using input, the number of called SEs was notably lower in samples with gene amplification, owing to very high H3K27ac signal in amplified regions, thus creating a bias in the ranked enhancer plot (Fig. S3A). To correct for the copy-number bias, we set a threshold and defined the top 1000 enhancer regions in each sample as SEs for all downstream analyses, approximately representing the mean number of SEs per sample (mean = 956 SEs, Table S2).

To gain insight into subtype-specific enhancer alterations and assess sample similarity based on these SE profiles, we determined the global repertoire of SEs in bladder by extracting a consensus set of 2887 SEs present in at least 2 of our 24 samples (Table S3). Using PCA of H3K27ac signal corresponding only to the consensus SE regions, we again found that samples were grouped according to molecular subgroup, separating Basals from Luminals, NMIBC segregating with differentiated tumors (Fig. 3A). This reveals that the variability in SE profiles reflects the differences in Basal and Luminal transcriptional programs. We also performed PCA using H3K27ac, H3K27me3 and H3K9me3 independently for peaks located inside the SE consensus regions for the tumour samples alone (Fig. S3B). Interestingly, all three profiles separated the Ba/Sq tumors from the other samples, indicating that all three histone marks are likely linked to SE regulation of bladder cancer molecular subtypes.

**Figure 3:**
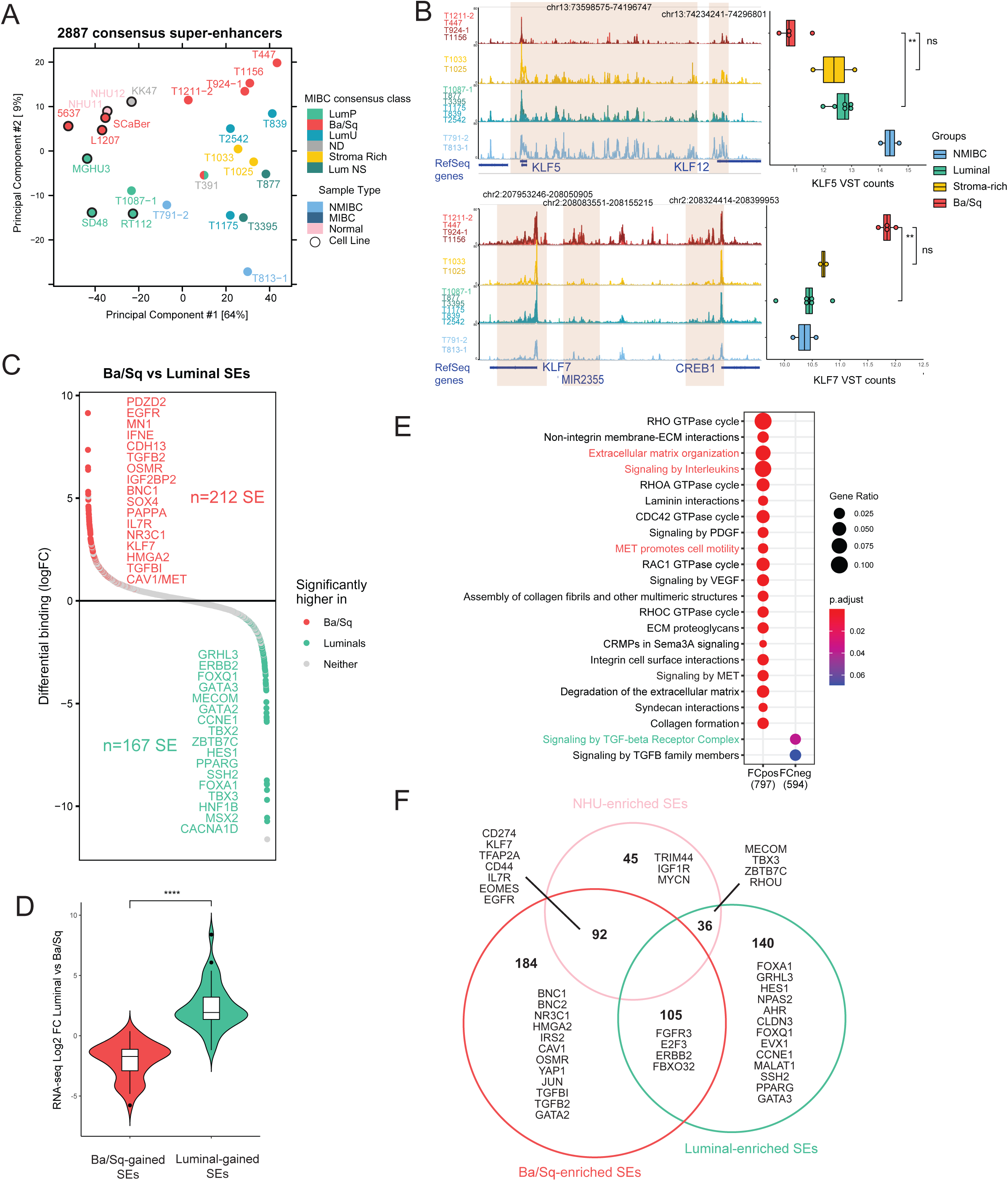
**Identification of the bladder super-enhancer repertoire and subtype specificities** A. PCA of H3K27ac signal inside ROSE consensus SE (n=2887) for all samples. B. Representative examples of H3K27ac signal in Ba/Sq, Stroma-Rich, Luminal and NMIBC tumors with corresponding RNA-seq gene expression. Orange boxes represent SE localisation. C. Fold Change plots for differentially bound SEs between Ba/Sq and Luminals samples. Significance by p-value <0.05. D. Plot comparing expression LogFC between Ba/Sq and Luminal samples for genes assigned to subgroup-enriched SE. E. Reactome pathway enrichment analysis of genes associated with Ba/Sq vs Luminal SEs. F. Venn diagram comparing 3 differential analyses of SE. NHU-enriched SEs are enriched in NHU vs Luminal or NHU vs Basal (pink circle). Basal-enriched SEs are enriched in Basal vs NHU or Basal vs Luminal (red circle). Luminal-enriched SEs are enriched in Luminal vs NHU or Luminal vs Basal (green circle).

We sought to further explore the functional pathway enrichment of SE-regulated genes. We first assigned SEs to their closest and most transcriptionally correlated genes (see methods). For example, a large SE mapped close to *KLF5*, whose expression was correlated with SE activity mostly in Luminal samples, as previously reported (31) (Fig. 3B). In contrast, *KLF7* was regulated by a SE mostly active in Basal samples (Fig. 3B). We then performed pairwise differential analyses between subtypes (Fig. 3C, S3C). The comparison between Basal and Luminal samples identified 369 subgroup-specific SEs (Fig. 3C, Table S3). By comparing the differential SEs to RNA-seq differential expression analysis, we confirmed that luminal-gained SEs showed significantly higher expression levels in Luminal samples relative to Basal samples and vice-versa for basal-gained SEs (Fig. 3D). We then validated the subgroup-specific SE-associated genes identified in our samples in a larger dataset, leveraging the gene expression profiles of the TCGA-BLCA MIBC cohort across molecular subtypes (n=406) (32, 33). Hierarchical clustering of TCGA samples using the genes associated to the most differentially regulated SEs in our consensus repertoire recovered the molecular classification (Fig. S3D, Table S3). Strikingly, differential analysis between Basal and Luminal SEs revealed that Luminal-specific SEs were attributed to known transcriptional drivers of the luminal phenotype, namely *GATA3*, *PPARg*, *FOXA1* (14, 22). Luminal-gained SEs were associated with “Signalling by TGF-beta family members”, notably due to SEs annotated close to negative regulators of TGF-beta signalling such as the E3 Ubiquitin ligase *SMURF1* or *SMAD6 (*Fig. 3C, E, Table S4). In contrast, SE regions significantly bound at higher levels in the Basal tumours were associated with genes known to contribute to Basal cancer biology such as *EGFR,* but also less characterized genes with regards to bladder cancer biology, such as genes related to inflammation and FOXO signalling (*IL7R*, *FBX032*), signalling by Interleukin or signalling by MET, the activation of which is often correlated with BLCA progression (34) (Fig. 3E). We also identified genes encoding membrane receptors (*IL7R*, *OSMR, EGFR*) and transcriptional regulators (*BNC2*, *HMGA2*, *KLF7*, *NR3C1*) as enriched in Basal tumours (Fig. 3C). Taking advantage of the NHU samples in our cohort, we extracted differential SEs in three comparisons (Ba/Sq *vs* NHU *vs* Luminals, Fig. S3C, Fig. 3F). This analysis validated the identification of genes that could be specific to cancer biology, such as *IL7R, OSMR, JUN, NR3C1* in Ba/Sq subtype, or *NPAS2*, *FOXQ1*, *GRHL3* in Luminal samples (Fig. 3F). Overall, we established a first SE repertoire for bladder cancer, highlighting subgroup-specific, cancer-specific SE activation coupled with gene expression.

### Super-enhancers regulate a network of candidate master transcription factors for bladder cancer subgroups

SEs often regulate the expression of master TFs, forming autoregulatory loops and correlated networks (35, 36). Having established the SE landscape in bladder cancer, we next sought to determine which master regulators control the subtype-specific transcriptional programs. To this end, we overlaid the genomic coordinates of subgroup-specific peaks inside SEs with publicly available ChIP-seq datasets (37, 38). Our analysis revealed that Luminal-specific SEs were significantly enriched in several TF binding sites (Fig. 4A, Table S5), including known regulators of Luminal subtypes FOXA1, GATA3, and ESR1 (3, 14, 32). Basal-specific SEs were enriched in binding sites of a different set of regulators, including components of the AP-1 complex (FOSL1, FOSL2, JUND, JUNB), as well as SMAD2/3, NFkB, and STAT3. Further DNA motif enrichment analysis comparing Basal differential peaks inside subgroup-specific SEs over Luminal ones, again identified AP-1 as a potential regulator of Basal SEs, as well as FOXA1, FOXA1:AR, GATA, and GRHL1/2/3 for Luminal SEs (Fig. 4B, Table S5, Homer (39)).

**Figure 4:**
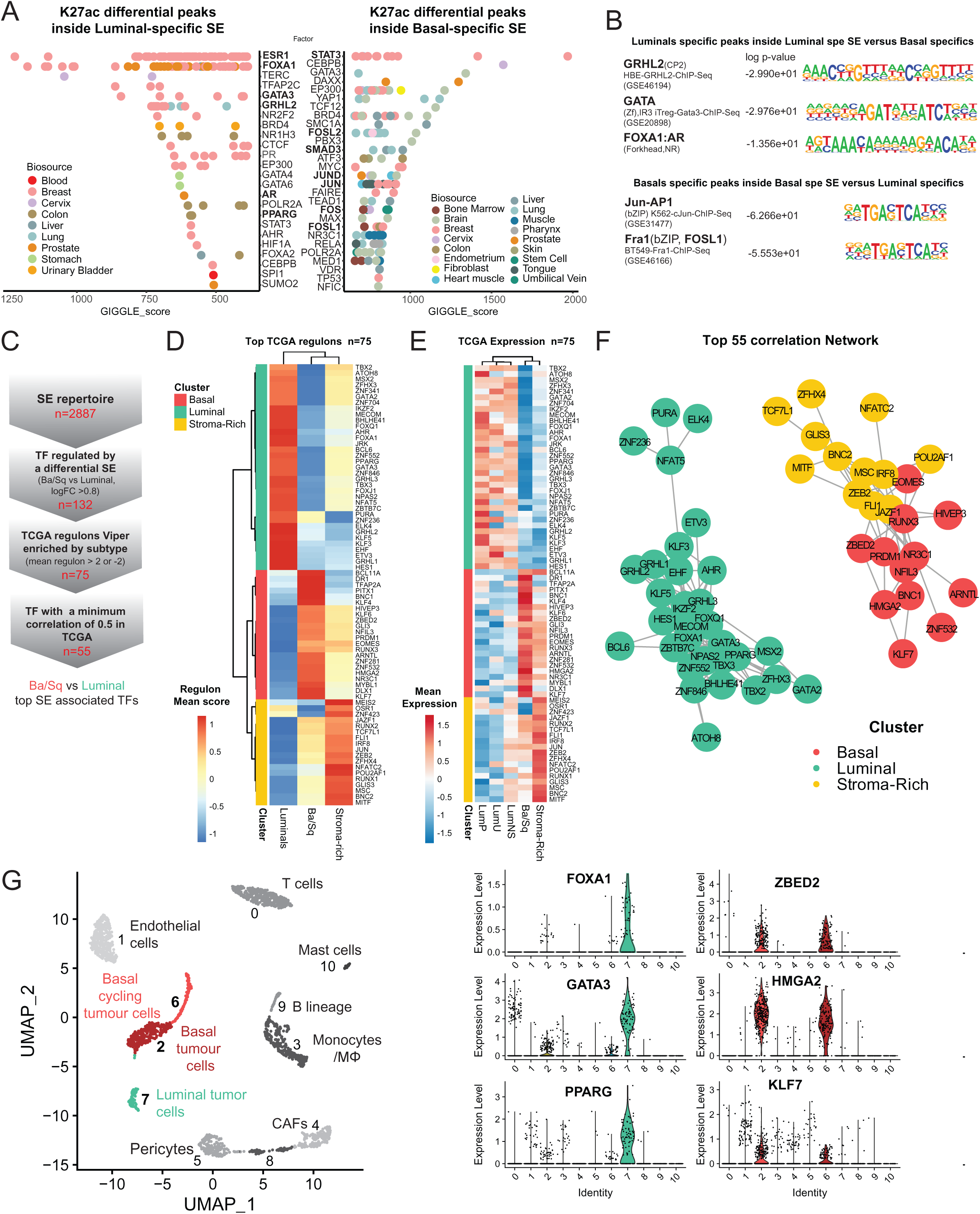
**Super-enhancers regulate a network of candidate master transcription factors for bladder cancer subgroups** A. Cistrome analysis of LumP and Ba/Sq specific SE. B. Homer motif enrichment analysis in H3K27ac differential peaks inside differential SE in Luminal vs Basal and Basal vs Luminal. C. Methodology to identify key coregulated SE-associated TFs. D. Heatmap of the top 75 TFs with high regulon score. Clustering identified 3 major clusters. E. Heatmap of the top 75 TFs expression in TCGA-BLCA. F. Correlation network of the top 55 TFs with an expression correlation coefficient of min 0.5 in TCGA-BLCA cohort. G. Single cell RNA-seq analysis of one Bladder Cancer tumors with both Basal and Luminal Population (GSM4307111) Right panel, associated expression for key TF in each compartment.

However, motif binding and ChIP-seq data are not available for all known TFs. To overcome this issue, we designed a method to identify subgroup-associated TFs and their co-regulated networks based on our differential SEs and the large transcriptomic cohort from the TCGA (32) (Fig. 4C). We selected TFs that were regulated by differential SEs (Basal vs Luminal), according to annotations from Lambert *et al*. (40), and that were differentially expressed in TCGA Ba/Sq vs Luminal subgroups. Then, we used ARACNe (41) and VIPER (42) analysis to identify and evaluate the regulons (group of genes regulated in response to one transcription regulator) of the 75 resulting TFs. Hierarchical clustering of the resulting TF regulons clustered scores into three groups, which were respectively associated with Luminal, Ba/Sq or Stroma-Rich subtypes (Fig. 4D, E). Since master TFs form interconnected networks with highly correlated levels of expression, we selected only TFs whose expression was correlated with that of at least one of the other TF in TCGA-BLCA data (Pearson correlation coefficient ≥0.5, n=55), and built the top correlated network based on subtype-specific SE-associated TFs (Fig. 4F) (see methods). This strategy identified a large module of Luminal TFs, including known Luminal-associated TFs (e.g., FOXA1/GATA3/PPARG), as well as TFs with yet unexplored roles in Luminal bladder cancer biology (e.g., HES1, FOXQ1, ZBTB7C, MECOM, GRHL2/3 and TBX3). Unlike for Luminals, few TFs have been characterized as key regulators of Basal tumours. Our analysis revealed a network of TFs whose activity could be essential to Basal bladder cancer biology, including HMGA2, KLF7, NR3C1 and ZBED2. Notably, ZBED2 has recently been associated with basal identity in keratinocytes (43) and regulation of inflammation in pancreatic cancer (44). Combining analyses of tumour and cell lines SEs should avoid the identification of master TFs expressed by the stroma. In fact, we found that TFs associated with the Luminal network showed strong expression correlation in TCGA-BLCA and in CCLE bladder cell cohorts (45) (Fig. S4A, B) while expression correlations of Stroma-Rich or Basal-associated TFs (e.g., ZEB1, SPI1) in the TCGA were lower for urothelial cell lines in the CCLE (46) (Fig. S4A, B). This indicates that expression of those TFs could be dependent on growing conditions and/or interactions with the tumour microenvironment.

To validate our Luminal- and Basal-specific TF networks, we analysed public single-cell RNA-seq data of a tumour presenting both a Luminal and a Basal cell population (GSM4307111, Fig. S4C). The Luminal-associated TFs FOXA1, GATA3 and PPARG were mostly expressed in the Luminal cell cluster, whereas ZBED2, HMGA2, and KLF7, newly identified as part of the Basal TF network, were mostly expressed in the Basal cell clusters (Fig. 4G), validating our subgroup-specific networks. Together, these analyses identified a targeted subset of interconnected candidate master TFs that could represent key regulators of bladder cancer subgroup identity.

### FOXA1 binds subgroup-specific bladder super-enhancers and correlates with their activation

We identified FOXA1 as one of our candidate master TFs for the Luminal bladder cancer subgroup. FOXA1 is a known pioneer factor with a demonstrated impact on Luminal bladder cancer biology (29, 32, 47), though the mechanism for how FOXA1 regulates cell identity is unknown. To better assess the role of FOXA1 in the regulation of bladder cancer SEs, we mapped FOXA1, CTCF (Insulator/enhancers) and H3K4me3 (Promoter) binding by ChIP-seq in two bladder cancer cell lines: SD48 (LumP) and 5637 (Ba/Sq). FOXA1 binding was mostly found outside promoters (Fig. S5A), with 61,083 FOXA1 peaks detected in SD48 cells and 39,445 in 5637, an expected variation as FOXA1 was more abundant in Luminal cells. Despite such differences, we identified three classes of FOXA1 peaks: SD48-specific peaks, peaks overlapping in the two cell lines, and 5637-specific peaks (Fig. 5A, Fig. S5B), which suggests that FOXA1 has specific targets in each cell line and subtype. Interestingly, when analysing TF binding sites from publicly available ChIP-seq data, SD48-specific FOXA1 peaks were highly enriched not only for FOXA1 binding sites, but also GATA3 binding sites (Fig. 5B), which could indicate a functional partnership between FOXA1 and GATA3 for regulation of the Luminal program, as suggested by Warrick *et al.* and described in breast cancer (14, 48). Surprisingly, 5637-specific FOXA1 peaks were mostly enriched at AP-1 binding sites and not FOXA1 sites (Fig. 5B, Fig. S5B). Both enrichments were confirmed by Homer motif analysis of SD48-specific peaks versus 5637-specific peaks or vice and versa (Fig. S5C and D, (39)). Ontology comparison of genes associated with the three classes of FOXA1 peaks showed that 5637-specific peaks were enriched in terms associated with Ba/Sq super-enhancers (e.g. Signalling by Tyrosine Kinase, Signalling by MET, Signalling by Interleukin), indicating that FOXA1 might be involved in the regulation of both Luminal and Basal bladder cancer subtypes (Fig. 5C). Indeed, FOXA1 binding in the two cell lines overlapped with most (87%) of the total repertoire of bladder SEs (Fig. 5D) and correlated strongly with H3K27ac levels at these loci (Fig. 5E, F), in line with a role for its regulation of these SEs. Notably, FOXA1 bound at SEs associated with genes involved in regulating urothelial differentiation and strongly correlated with increased H3K27ac at these loci. This could clearly be observed for the Luminal-specific SEs associated with *GATA3* or *PPARG* in the Luminal SD48 cells and in both cell lines for the non-specific *PPARG* SE. But we also found FOXA1 binding associated with high H3K27ac at certain Basal-specific SEs in the Basal 5637 cells, such as *TGFB2* (Fig. 5G). This implies that FOXA1, even if expressed at a low level as in Ba/Sq cells, could play an important role in BLCA biology, through enhancer/SE regulation. In summary, FOXA1 may regulate bladder cell identity through binding of subgroup-specific bladder SEs with partners such as GATA3 in Luminal cells and AP-1 in Basal cells.

**Figure 5:**
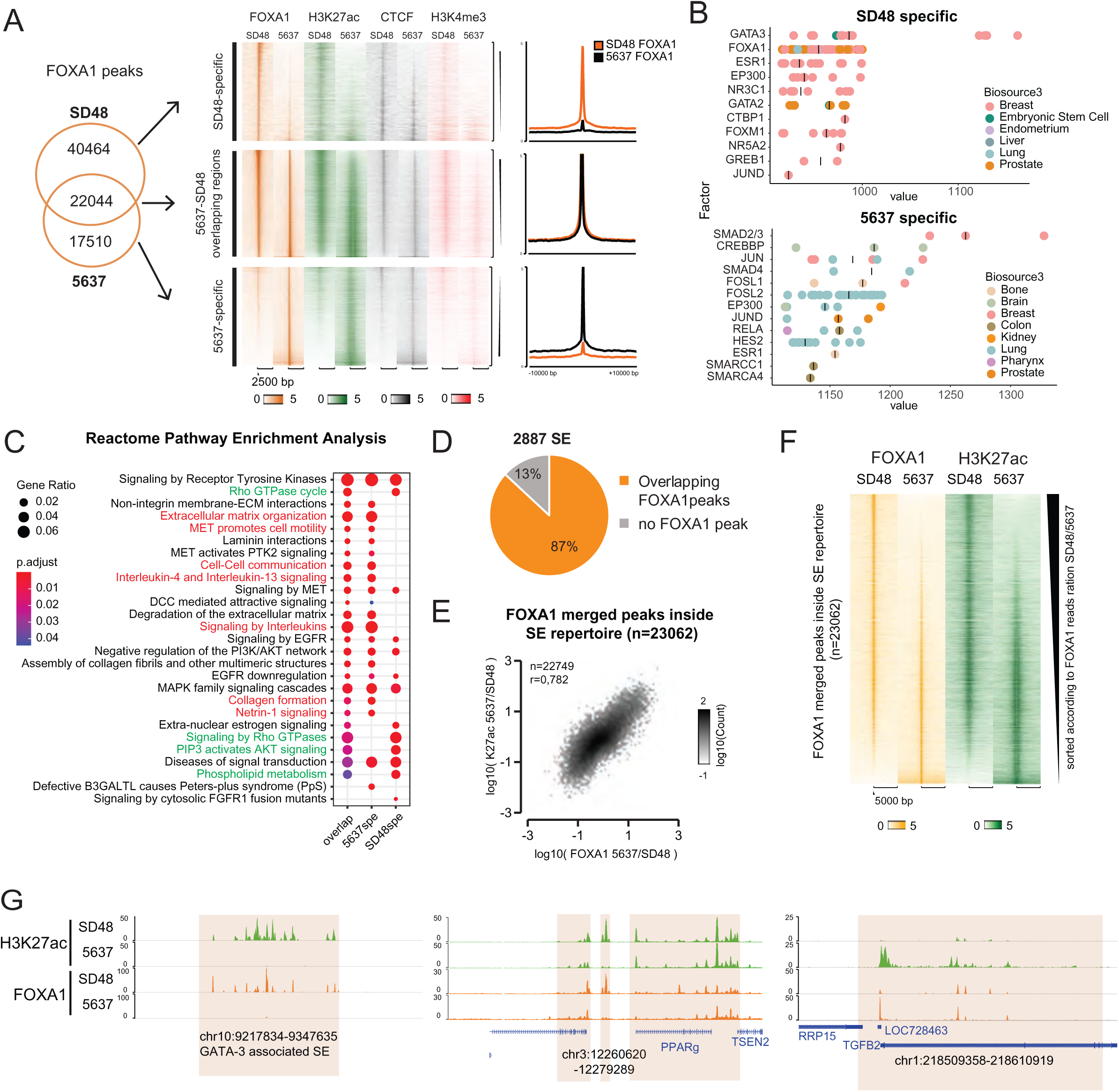
**FOXA1 binds subgroup-specific bladder super-enhancers and correlates with their activation** A. Venn diagram comparing FOXA1 ChIP-seq peaks in SD48 and 5637 cell line. Heatmap for 3 categories of peaks and associated mean profiles. B. Cistrome analysis of motif enrichment analysis in SD48-specific and 5637-specific FOXA1 peaks. C. Reactome pathway analysis of genes associated to the 3 categories of FOXA1 peaks. D. Pie chart showing proportion of SE with an overlapping FOXA1 peak (merge of FOXA1 peaks in SD48 and 5637). E. Correlation between H3K27ac peaks versus FOXA1 peaks inside SE. F. Heatmap of FOXA1 and H3K27ac reads on FOXA1 peaks overlapping Super-enhancers ranked by FOXA1 reads ration in SD48 vs 5637. G. Genome browser view of GATA3, PPARg and TGFB2 associated SE. SE are highlighted with orange boxes.

### FOXA1 regulates inflammation and cellular identity

To better understand FOXA1 function, we performed short-term (<72h) knock-down in both Luminal and Basal models. Knock-down of FOXA1 by siRNA decreased clonogenicity and proliferation of both Luminal and Basal cells (Fig. S6A, B) and it reduced cell viability in both RT112 (LumP subtype) and SCaBER (Ba/Sq) cells, with a stronger impact in RT112 (Fig. 6A). Furthermore, FOXA1 knock-down in RT112 and SCaBER cell lines dramatically altered gene expression (Fig. 6B, Table S6). The downregulated genes were related to cell cycle and checkpoint pathways, consistent with the reductions in viability and proliferation upon FOXA1 knock-down. Surprisingly, the upregulated genes in both cell lines were strongly associated with inflammatory signalling and interferon response (Fig. 6C, Fig. S6D). Notably, FOXA1 knock-down induced upregulation of master interferon response TFs, STAT1 and STAT2, and key genes involved in the regulation of inflammation in human cancer, including the immune checkpoint modulator *CD274* (PD-L1) (Fig. 6D), which we also identified as a downregulated SE in both Luminal and Basal *vs* NHU cells (see earlier Fig. 3F). While our FOXA1 ChIP-seq in Luminal and Basal cell lines showed FOXA1 binding at many interferon responsive genes, we did not observe strong FOXA1 enrichment on *STAT1, STAT2* or *CD274* promoters or enhancers (Fig. S6E). This suggests that the upregulation of these genes upon FOXA1 knock-down is independent of FOXA1 binding of their regulatory elements, in agreement with recent work showing that FOXA1 directly binds and inhibits the STAT2 protein to dampen inflammation in a chromatin-independent manner (49). Interestingly, if FOXA1 knock down triggered interferon response in both Luminal and Basal models, its depletion affected the Luminal network of co-regulated TFs only in RT112 cells and not in SCaBER (Fig. 6E). PCA projection of TCGA-BLCA transcriptomes together with that of our knock down cells on our scRNA-seq-derived Basal/Luminal signature space confirmed that FOXA1 acute depletion induced a small but consistent shift from Luminal towards Basal subtype only in RT112 cells (Fig. S6F, see methods). Therefore, in agreement with a previous study (14), short-term knock-down of FOXA1 showed a consistent but mild impact on cell identity, not sufficient to majorly alter the subtype of the luminal cells.

**Figure 6:**
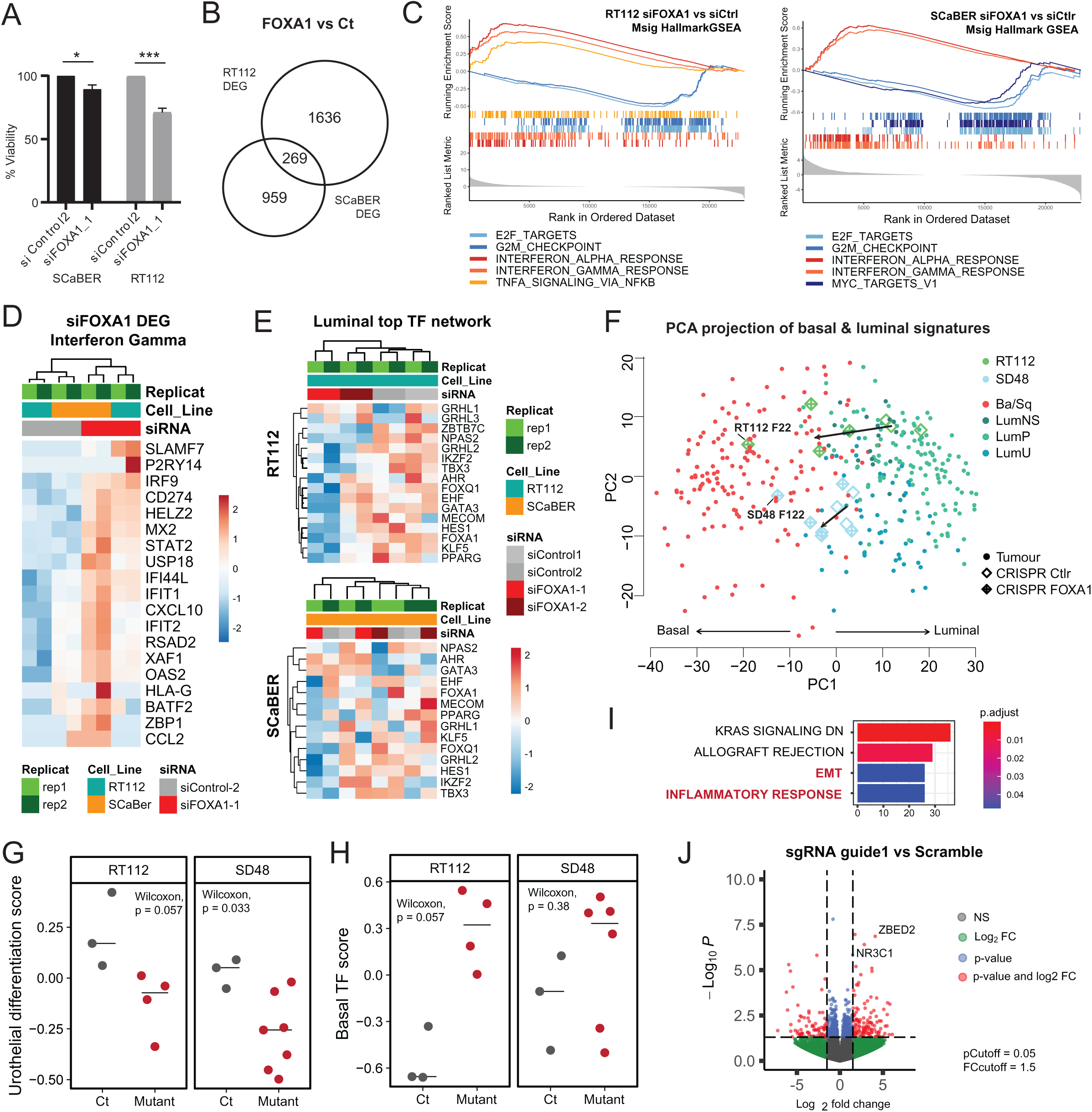
**FOXA1 regulates inflammation and cellular identity** A. Cell viability in RT112 and SCaBER under siRNA treatment against FOXA1 B. Venn diagram comparing differentially expressed genes in RT112 and SCaBER FOXA1 KD. C. GSEA plot of Msig Hallmark GSEA Analysis of genes differentially regulated in RT112 and SCaBER cell lines upon FOXA1 siRNA (2 independent siRNA, 2 replicates) D. Heatmap of genes in Hallmark interferon gamma response genes that are differentially regulated in FOXA1 KD *vs* Ct (min Fold Change = 1,5). E. Heatmap of Top Luminal TFs expression in RT112 and SCaBER cell lines upon FOXA1 KD. F. PCA projection of TCGA tumors and CRispR mutant clones on the Basal/Luminal signa-tures. G. GSVA analysis of FOXA1 CRispR mutant clones on Urothelial differentiation signature from Eriksson et al. H. GSVA analysis of FOXA1 CRispR mutant clones on Basal TFs identified in Fig. 4F I. Overrepresentation analysis of DEG in FOXA1 mutant *vs* Controls. J. Volcano plot of Deseq2 RNA-seq analysis comparing pooled CRispR mutant FOXA1 clones in SD48 and RT112 *versus* controls.

Altering the epigenetic landscape could indeed take a longer time. To determine if FOXA1, through its binding to the SE repertoire, regulates the bladder cancer epigenetic landscape and subsequently cellular identity, we produced *FOXA1* CRISPR mutant clones allowing long-term FOXA1 inactivation in two Luminal cell lines (SD48 and RT112, Fig. S6G). Despite fundamental differences between RT112 and SD48 cellular models and heterogeneity between clones due to clonal selection, transcriptomic analysis of 3’ RNA-seq data by PCA distinguished CRISPR *FOXA1* mutant clones from wildtype (WT) (Fig. S6H). Importantly, PCA projection of TCGA-BLCA transcriptomes together with that of our WT and mutant clones on the Basal/Luminal signature space showed that mutation of *FOXA1* induced a strong shift from the Luminal cluster to the Basal cluster (Fig. 6F). GSEA analysis confirmed that *FOXA1* mutants were enriched for our Basal signature and depleted for our Luminal signature (Fig. S6I)(26, 27). GSVA analysis further revealed that *FOXA1* mutant clones were less differentiated than WT controls (Fig. 6G, (11)) and tended to express higher levels of TFs associated with the Basal TF network (Fig. 6H, Fig. 4F). Differential gene expression analysis revealed 1040 and 1102 Differentially Expressed Genes (DEGs) in RT112 and SD48, respectively, when comparing *FOXA1* mutant clones to WT (Fig. S6J). *FOXA1* mutant DEGs were associated with EMT, KRAS signalling and the inflammatory response pathway (Fig. 6I), all linked to Basal phenotypes. Intriguingly, differential analysis of *FOXA1* mutants vs WT revealed increased expression of NR3C1 and ZBED2 in the mutants, two of the candidate master TFs identified in our Basal TF network (Fig. 6J, Table S7). In summary, our results demonstrate that loss of *FOXA1* promotes a clear shift from Luminal to Basal cell identity.

### ZBED2, a novel Basal-associated TF involved in inflammation dampening

To further explore the interconnected network of candidate master TFs, we chose to examine ZBED2 as one of the TFs in the Basal network since it was upregulated by *FOXA1* CRispR inactivation, and because of recent work in keratinocytes that identified a role for ZBED2 in the basal phenotype (43). ZBED2 expression in the TCGA-BLCA cohort is upregulated in the Ba/Sq subtype (Fig. 7A) and correlates with poor survival prognosis (Fig. S7A). ZBED2 expression is negatively correlated with FOXA1 expression in the TCGA cohort (Fig. 7B), but more interestingly scRNA-seq in CCLE bladder cell lines shows that FOXA1 and ZBED2 expression are often mutually exclusive (Fig. 7C). As little is known about the ZBED2 TF, we used ARACNE/VIPER algorithms to identify the ZBED2 regulon based on TCGA-BLCA expression data. Interestingly, FOXA1 was predicted as a ZBED2 target, with the most negative weight, whereas two genes associated with Basal-specific SEs (*IL7R* and *CAV1*) were in the top 10 positive ZBED2 regulon weights (Fig. S7B). Using ZBED2 ChIP-seq data from pancreatic cancer cell lines (44) (the only ZBED2 ChIP-seq reported so far), we found a high confidence ZBED2 peak in the *FOXA1* promoter (Fig. 7D, left). Analysis of the RNA-seq data from the same study revealed that ZBED2 overexpression triggered downregulation of FOXA1 (Fig. S7C, *p*=0.004). In our data, we found that the *ZBED2* SE was highly enriched in FOXA1 binding in SD48 luminal cells, whereas FOXA1 binding was significantly decreased in 5637 Ba/Sq cells, and negatively correlated with ZBED2 expression (Fig. 7D, right). Overall, these findings suggest that FOXA1 and ZBED2 could negatively regulate each other to promote or maintain Luminal or Basal identity, respectively.

**Figure 7:**
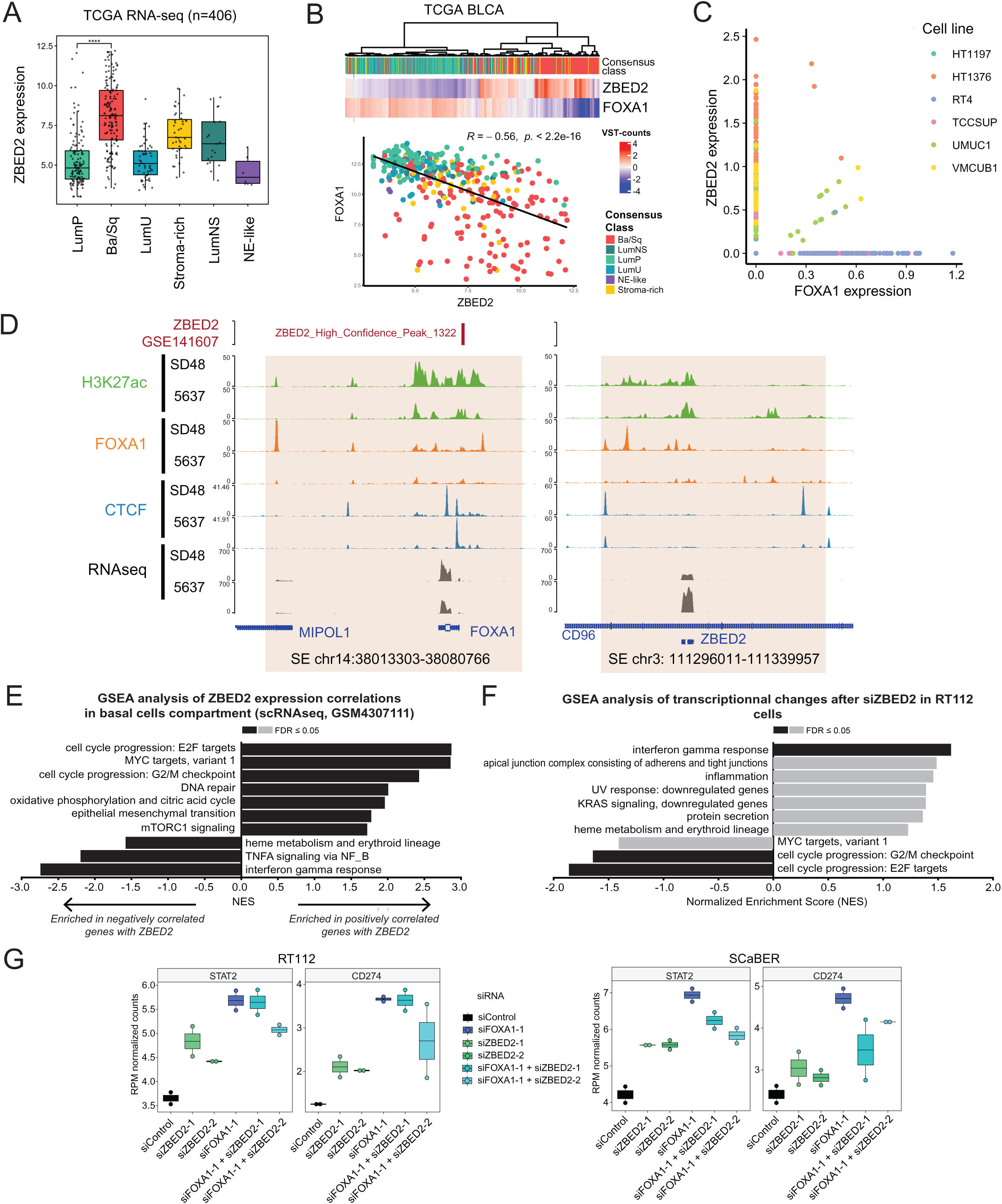
**ZBED2, a novel Basal-associated TF also involved inflammation dampening** A. TCGA expression of ZBED2 by Subtypes. B. TCGA expression Heatmap of ZBED2 and FOXA1 and TCGA correlation between ZBED2 and FOXA1. C. Expression of FOXA1 and ZBED2 in single-cell transcriptomics from bladder cancer cell lines in the Cancer Cell Line Encyclopedia (CCLE), highlighting the nearly mutually exclusive expression of these genes. D. Genome browser view of ZBED2 and FOXA1 loci in SD48 and 5637 cell lines E. GSEA analysis (Hallmarlk) of ZBED2 correated genes in basal cells population of GSM4307111 scRNAseq tumor. F. GSEA analysis (Hallmarlk) of gene expression upon siZBED2 KD in RT112 (siZBED2-1 and siZBED2-2). G. 3’seq STAT2 and CD274 (PD-L1) expression in RT112 and SCaBER after siZBED2 and siFOXA1.

On the other hand, ZBED2 has been shown to inhibit STAT2 and dampen inflammation by direct competition with IRF1 for Interferon Responsive Element binding in the pancreas (44). We therefore sought to determine if ZBED2 is involved in the downregulation of interferon signalling in bladder cancer, potentially through interfering with FOXA1-activated pathways. Intriguingly, *ZBED2* expression in the TCGA-BLCA cohort positively correlated with interferon gamma associated gene expression (Fig. S7D), which could be an indication that ZBED2 increases in response to inflammation at the cell population level, or vice versa. We then examined the correlation between ZBED2 expression and different cellular pathways at the single-cell level using publicly available scRNA-seq data (50). Our analysis revealed that *ZBED2* expression anti-correlates with interferon response and positively correlates with cell cycle progression and *E2F* targets within the same cell (Fig. 7E), suggesting that the positive correlation with interferon response in the bulk RNA-seq data reflects increased levels of *ZBED2* expression and interferon response genes in different subpopulations of cells. To test this association further, we knocked down *ZBED2* by siRNA and performed bulk 3’ RNA-seq in two BLCA cell lines. Strikingly, downregulation of *ZBED2* increased expression of interferon response genes and decreased expression of cell cycle progression and *E2F* target genes (Fig. 7F). Furthermore, siZBED2 in both RT112 and SCaBER cells increased gene expression of *STAT2* and *CD274* (Fig. 7G) and tended towards decreased cell viability (Fig. S7E). Therefore, ZBED2 directly dampens interferon response in bladder cancer, in agreement with its reported role in the pancreas (44). Notably, siRNA of *FOXA1* induced strong *STAT2* and *CD274* expression, while double knock-down of both *FOXA1* and *ZBED2* partially dampened this response compared to siFOXA1 alone (Fig. 7G), suggesting that the inflammatory response resulting from FOXA1 knock-down is partially dependant on ZBED2 target genes. In conclusion, both FOXA1 and ZBED2 inhibit inflammatory response and promote bladder cancer cell survival.

## Discussion

Epigenetic mechanisms are essential for the establishment and maintenance of cellular identity notably through SE regulation of master transcriptional regulators (17). Bladder cancer has been extensively studied at the transcriptomic level, but until two recent studies, very little was known about its epigenetic landscape (21).

Here, we report a large epigenetic profiling of both bladder cancer primary tumours and bladder cancer cell lines representative of the main molecular subtypes, as well as NHU cultures, using three histone marks ChIP-seq and paired RNA-seq. Using integrative analyses, we established a comprehensive chromatin state map of bladder cancer and showed that Basal and Luminal subgroups can be distinguished by their chromatin profiles. This map can be used to identify new genes or regulatory regions for diagnostic, prognostic or pharmacological targeting.

We characterized the bladder SE repertoire, and through differential analysis identified subgroup-specific and cancer-specific SE activation. Our study corroborates and expands the two recently reported enhancer landscapes of bladder cancer (20, 21). Consistent with prior reports, Luminal-activated SEs were located in proximity to known key regulators of the Luminal phenotype, namely *FOXA1, GATA3,* and *PPARG* (3, 14, 21), and to new Luminal-associated genes, such as *NPAS2 and GRHL2 –* also identified by Iyyanki *et al*. (20) *–* or *KLF5,* recently characterized as activated through super-enhancer amplification in various squamous cell carcinomas (31). Importantly, based on our data for 7 Ba/Sq samples, we were able to identify higher enhancer activity associated with potential key genes in Basal tumour biology, including cell surface receptors (*IL7R*, *OSMR*, *EGFR*, *MET*) and transcriptional regulators (*BNC2*, *HMGA2*, *KLF7*, *NR3C1*).

We further characterized subgroup-associated master regulators and co-regulated networks for both Luminal and Ba/Sq subgroups using two complementary approaches, in order to overcome issues linked to low expression, unknown binding motifs or multi-partner complexes. First, we identified TFs with enriched binding sites or DNA motifs in subgroup-specific SEs. Second, we combined SE activity in our cohort with regulon analysis of TCGA data to identify master regulator networks for Luminal and Basal subgroups. The first approach, based on public ChIP-seq data, validated the role of TFs involved in urothelial differentiation in Luminal SE activity, namely FOXA1 and GATA3, but also revealed that the AP-1 complex regulates Basal SEs. AP-1 has been shown to drive reprogramming of breast cancer cells from a Luminal to a Basal phenotype during treatment resistance acquisition through high-order assemblies of transcription factors (48). Thus, a role for AP-1 in driving Basal enhancers and cell identity in bladder cancer suggests AP-1 inhibitors as potential therapeutic options for this aggressive disease. Interestingly, our mapping of FOXA1 binding sites in two different cell models indicated that the pioneer factor binds most bladder-associated SEs, even if its DNA binding motif is mostly found in the Luminal-specific SEs. The mapping of FOXA1 binding also confirmed that FOXA1 binding sites in Basal enhancers are associated with AP-1 localisation, suggesting that AP-1 could play an important role in the regulation of Basal regulatory regions through FOXA1 recruitment—or trapping—at discrete chromatin loci. By combining the ChIP-seq approach with regulon analysis, we were able to highlight new Luminal-associated TFs, in addition to known Luminal master regulators (FOXA1, GATA3, PPARG) and the recently identified NPAS2 (20). Importantly, we also identified a Basal TF network, including ZBED2, KLF7, HMGA2, NR3C1 as major regulators, whose expression was restricted to the basal component of tumours, according to scRNA-seq data. To our knowledge, the role of these TFs has not been investigated in bladder cancer biology.

With regards to ZBED2, a scRNA-seq study revealed that it promotes basal cell identity of keratinocytes (43). Another study demonstrated that in pancreatic cancer, ZBED2 represses differentiation and dampens STAT2-mediated inflammatory response through IRES binding competition with IRF1 (44). Previous work showed that the three master Luminal TFs (FOXA1, GATA3, and PPARG) had to be perturbed simultaneously to induce a cell identity switch from luminal to basal (14). However, we found that while short term knock-down of FOXA1 had a mild effect on cell identity, the long-term inactivation of FOXA1 alone through CRISPR mutation was sufficient to induce a shift from Luminal to Basal subgroup in luminal cells, highlighting the role of FOXA1 in the regulation of cell fate. Moreover, we demonstrated that this major shift is accompanied by activation of one of our newly identified Basal network TFs, ZBED2. Despite its known role as an activator of transcription, FOXA1 has also been associated to direct repression of transcription (51). ZBED2 is described as a transcriptional repressor (44). Therefore, FOXA1 and ZBED2 could repress each other, defining a new cell identity regulatory loop. Through functional knock-down and knock-out experiments, we verified that FOXA1 and ZBED2 have antagonistic but interconnected functions in cell identity. However, ZBED2 is expressed at a very low level and additional experiments, including overexpression models are needed to validate its repressive function on FOXA1, or vice versa, and its potential role in Luminal to Basal plasticity.

Finally, our work also uncovers a role for both FOXA1 and ZBED2 in the regulation of inflammation in bladder cancer. While they play antagonistic roles in the regulation of cell identity, we found that they share a common function in inhibiting inflammation. Short term loss of either FOXA1 or ZBED2 triggers an inflammatory response, identified through STAT2 overexpression, in agreement with the study of ZBED2 function in the pancreas [44]. The low FOXA1 binding enrichment at STAT2 in our FOXA1 ChIP-Seq experiments suggest that FOXA1 could have a repressive function of inflammation presumably independent of its chromatin binding. These conclusions are in accordance with the recent work of He et *al.* [57] characterizing a chromatin independent function of FOXA1, which, by direct binding of STAT2 protein, inhibited STAT2-mediated inflammation. This could explain the limited infiltrate of luminal tumours, expressing high levels of FOXA1.

Therefore, given the dual role of FOXA1 and ZBED2 in the regulation of cell identity and inflammation, it will be important to study their link with tumour plasticity and in response to immunotherapy. Although direct inhibitors do not yet exist, targeting FOXA1/ZBED2 or the upstream or downstream signaling pathways, may improve sensitivity to immune-based therapies. Similarly, it will be worth studying the effect of interferon treatment on FOXA1 and ZBED2 inhibited inflammation as it could be used to overcome the inflammation inhibition induced by these two master regulators.

If FOXA1 and ZBED2 revealed promising features, our study identified numerous other super enhancers, associated genes and master regulators that could be explored for pharmacological targeting.

General targeting of SEs with BRD4 inhibitors has shown efficiency, in particular in cancers with specific SE single mutation alterations or with the activation of MYC SE in leukemia or lymphoma (18, 52, 53). However, those treatments show mild efficiency in solid tumours and enhancer rewiring has been associated to resistance to treatment. Identification of SEs associated with bladder cancer and subgroups may pave the way for further research into targeting activated master regulators, upstream/downstream activated pathways or even with the advent of RNA and CRISPR technology, directly targeting SEs.

## Conclusions

We provide an integrated epigenomic and transcriptomic map of bladder cancer constituting a new comprehensive tool to study epigenetic regulation of muscle-invasive bladder cancer. We revealed Luminal and Basal coregulated networks of super-enhancers and associated transcription factors as new potential targets with important clinical relevance. Our findings and functional assays on FOXA1 and ZBED2 demonstrate that the SEs and TF networks identified herein represent prime targets for further pre-clinical investigation for bladder cancer treatment.

## MATERIALS and METHODS

### Biological Resources

#### Cell lines and culture

The human bladder cancer derived cell lines RT112, 5637, KK47, and SCaBER were obtained from DSMZ (Heidelberg, Germany). MGH-U3, KK47 and SD48 cell lines were provided by Yves Fradet (CRC, Quebec), Jennifer Southgate laboratory (previously of Cancer Research Unit, St James’s University Hospital, Leeds, UK), and Henri Mondor Hospital (Créteil, France), respectively. The L1207 cell line was derived from tumour T1207 (25). RT112 and 5637 cells were cultured in RPMI medium, L1207 were cultured in DMEM-F12 and all the other cell lines were cultured in DMEM medium. All cell media were supplemented with 10% fetal bovine serum (FBS). Normal human urothelium (NHU) cells were obtained from normal ureter urothelium from healthy kidney donors from Foch hospital and were cultured as previously described (26). All cells were cultured at 37°C in an atmosphere of 5% CO2 and were routinely tested for mycoplasma contamination.

#### Patient tumour tissue processing

We selected human tumours with an available OCT-compound frozen block from our CIT (Carte d’Identité des Tumeurs) cohort (9, 22). Each block was frozen-sectioned and stained with hematoxylin and eosin. Pathology review was performed to confirm the tumour stage and to select tumour areas, in order to enhance neoplastic content (estimated at 30 to 95%, median tumour cell content = 65%). For tumours with sufficient material, tumour-enriched areas were macrodissected from the frozen block and manually finely ground in a mortar. Frozen ground tumour tissue was kept at -80°C until further processing. The characteristics of the tumours are shown in Table 1.

#### CRispR vectors, siRNA

siRNA and CrispR vectors used in the study are referenced in supplemental information file.

### Chromatin immunoprecipitation and sequencing

#### Tumour chromatin cross-linking and extraction

In order to obtain efficiently disrupted tissue, the frozen ground material (15mg) was further homogenized using a tube pestle or the TissueLyser II system (Qiagen). Disrupted tissue was then processed using the reagents from the iDeal ChIP-seq Kit for Histones (Diagenode), according to the manufacturer’s instructions. Briefly, the tissue was homogenized and washed in 1ml PBS-protease inhibitor cocktail. DNA-protein cross-linking was ensured with an 8-minute incubation in 1% formaldehyde then quenched with 0.125 M glycine for 5 minutes. Cells were then washed and lysed. Centrifuged cell lysates were resuspended in shearing buffer and sonicated using the Pico Bioruptor device (Diagenode) for 15 minutes (30s ON/30s OFF). Following a centrifugation at 16000g for 10 minutes, an aliquot was reserved to control the sonication and the remaining supernatant was stored at -80°C. Sonication efficiency was controlled for each sample on the aliquot of sheared chromatin by overnight reverse cross-linking, DNA was purified using the phenol-chloroform method and 2% agarose gel electrophoresis was used to determine DNA fragment size.

#### Tumours ChIP-seq

Tumour samples with optimal chromatin fragment size (200-500 bp) were immunoprecipitated using the iDeal ChIP-seq Kit for Histones (Diagenode). Magnetic immunoprecipitation of sheared DNA-chromatin complexes (500ng) was performed overnight using a rabbit polyclonal histone H3K27acetyl ChIP Grade antibody (ab4729, Abcam), H3K27me3 (Active Motif, ref. 39155), and H3K9me3 (Active Motif, ref. 39161). Magnetic immunoprecipitation beads were washed the following day. The captured chromatin as well as non-immunoprecipitated input chromatin underwent elution and reverse cross-linking steps. DNA purification was performed using iPure magnetic beads. Immunoprecipitation (IP) efficiency was verified by qPCR according to the manufacturer’s protocol. Library preparation from IP DNA and input DNA was performed using the Diagenode MicroPlex Library Preparation kit v2. The resulting amplified libraries were assessed using the Bioanalyzer system 2100 (Agilent) and sequenced using the HiSeq 4000 platform (Illumina) as single-read 50 base reads, following Illumina’s instructions. Reads were aligned to the reference genome (Hg19) using Bowtie 1.0.0.

#### Cell line ChIP-seq

Cell lines cultures were crosslinked directly in the growing medium with formaldehyde 1% for 10 minutes at room temperature. The reaction was stopped by adding Glycine with a final concentration of 0.125M for 10 minutes at room temperature. Fixed cells were rinsed 3 times with PBS containing protease inhibitors, pelleted, and resuspended in lysis buffer (10mM EDTA, pH8, 50mM Tris-HCl pH8, SDS 1%). After centrifugation, the ChIP was performed using ChIP-IT High Sensitivity kit (Active Motif, Carlsbad, CA, USA), following the manufacturer’s instructions. Chromatin was sonicated in a bioruptor Pico device (Diagenode) for 10 min (30s ON/30s OFF). Sheared chromatin was immunoprecipitated using an H3K27ac antibody (Abcam ab4729). Sheared chromatin was used as input-DNA control.

ChIP-seq libraries were prepared using NEXTflex ChIP-Seq Kit (#5143-02, Bioo Scientific) following the manufacturer’s protocol (V12.10) with some modifications. Briefly, 10 ng of ChIP enriched DNA were end-repaired using T4 DNA polymerase, Klenow DNA polymerase and T4 PNK, then size selected and cleaned-up using Agencourt AMPure XP beads (#A63881, Beckman). A single ‘A’ nucleotide was added to the 3’ ends of the blunt DNA fragments with a Klenow fragment (3’ to 5’exo minus). The ends of the DNA fragments were ligated to double stranded barcoded DNA adapters (NEXTflex ChIP-Seq Barcodes - 6, #514120, Bioo Scientific) using T4 DNA Ligase. The ligated products were enriched by PCR and cleaned-up using Agencourt AMPure XP beads. Prior to sequencing, DNA libraries were checked for quality and quantified using a 2100 Bioanalyzer (Agilent). The libraries were sequenced on the Illumina Hi-Seq 2500 as single-end 50 base reads following Illumina’s instructions. Sequence reads were mapped to reference genome hg19 using Bowtie 1.0.0.

### ChIP-seq data analysis and integration

Peak detection was performed using MACS2 (model-based analysis for ChIP-seq v2.1.0.20140616) software under settings where input samples were used as a negative control. We used a default cutoff and -B option for broad peaks.

To identify enhancer regions in each tumour we used ROSE (Ranked Ordering of Super-Enhancers) algorithm (18, 30), with the following parameters: 12.5kb stitching distance, exclusion of promoter regions 2500 bp around TSS. For each sample, stitched enhancer regions are normalized, ranked and plotted. The regions above the inflexion point are considered super-enhancers by the algorithm. However, the number of called super-enhancers was notably lower in cases with a known amplified gene. The very high H3K27ac signal in the amplified region likely created a bias in the plot of ranked enhancers. To correct for this bias, we set a threshold to the top 1000 ranked enhancers to select candidate super-enhancer regions.

Heatmaps and PCA of ChIP-seq signal were performed using Diffbind R package (version 2.16.0) or Easeq (54). For super-enhancers analysis, the top 1000 SE regions of either tumours or cell lines were merged for a consensus using Diffbind. Then, H3K27ac signal was calculated in the consensus peak for each sample. Differential analysis between molecular subtypes was performed with Diffbind and DESeq2 default parameters using both IP and input bam files, and a file containing the consensus super enhancer regions evaluated for differential analysis as input. Regions with an pval <0.05 were considered differentially bound.

Genomic annotation and pathway enrichment analyses were performed using ChIPseeker, clusterProfiler and GREAT (28).

#### Chromatin Binding enrichment analysis

Factor binding analyses were performed using public data available in Cistrome DB Toolkit (37, 38). DNA binding motif analysis was performed using HOMER known motif function (39).

#### Genomic annotation of the SE regions and cis-regulatory genes

SE activity and gene expression was jointly analysed to determine the cis-regulatory between the SE and genes on proximity. In brief, the overlap / proximal genes corresponding to each SE were annotated using GREAT/ROSE tools, as the candidate proximal genes regulated by the SE (30, 55). The spearman correlation coefficients between SE activity (H3K27ac read counts, log2RPKM normalized) and the expression of the candidate genes (RLE normalized) were calculated in the tumours. The gene whose expression showing the highest correlation with the activity of the corresponding SE was determined as the gene most likely regulated by the SE. The SE-gene relationships within the top 1% were also given, not limited to the proximal genes. The number of germline single nucleotide polymorphisms (SNPs) within a given SE as well as their association with bladder cancer (median –log10 p-value) was provided based on the UK biobank GWAS summary statistics (Neale lab Round 2, ukb-d-C67, extracted from the MRC IEU OpenGWAS database) (56). The germline SNPs falling within the SEs and with published GWAS-level association with BCa or with –log10P.value > 5 in GWAS summary statistics (PhenoScanner v2 database) were provided as GWAS SNPs within the SEs (57). For the genes most likely regulated by a given SE, we provided their median CERES dependency score of all and urothelial cancer cells from the Cancer Dependency Map database (45), as well as the p-value for difference between the two. We checked if any bias compared to the background in mutation type (missense, non-sense, synonymous, etc.) for the protein coding genes by Chi-square test. We checked if they were within the list of established cancer genes, including the COSMIC Cancer Gene Census and Network of Cancer Genes 6 (58, 59).

#### Chromatin state analysis and correlation with expression

ChromHMM was used to identify chromatin states. The genome was analyzed at 1000 bp intervals and the tool was used to learn models from the 3 histone marks ChIP-seq reads files and corresponding Input controls. A model of 6 states was selected and applied on all samples. The 6 states identified were then given functional annotation based on histone marks enrichment and ENCODE published chromatin states.

We checked the genome-wide association between gene expression and chromatin states of the TSS, in both tumour and cell line samples. In each tumour / cell line, we classified the genes according to the chromatin states of the TSS. For genes with multiple TSS, the chromatin states showing frequency dominance was considered. We then calculated the median expression of the genes by their TSS categories in each sample, and assessed the distribution of the median expression by chromatin state across all tumour and cell line samples.

ChromHMM output files were concatenated using the unionbed function from BEDTools, by which a consensus sample-by-states matrix was created, where in each cell the chromatin state corresponding to the column’s chromosome region in the row’s sample, excluding regions from sexual chromosomes, with all samples included (n = 24, including 15 tumours and 9 cell lines).

We next performed unsupervised analysis of the integrated chromatin states in tumour and cell line samples.

#### Selection of most informative features

We first looked for the most informative features in the consensus sample-by-states matrix where in each cell the chromatin state corresponding to the column’s chromosome region in the row’s sample, excluding regions from sexual chromosomes, with all samples included (n = 24, including 15 tumours and 9 cell lines). We excluded genome regions of ’no mark’ state to enrich our feature selection with active regions and filtered features with top 1% Shannon’s entropy. Then to further select informative feature, we signed-rank transformed the data: For each state, given the constitution of the histone marks and association with gene expression, we performed a numeric transformation of the categorical states by assigning numeric values to categorical states, as 3 to E2 (Active Enhancer / Promoter), -3 to E4 (Repressed Chromatin), 2 to E3 (Bivalent Enhancer / Promoter), -2 to E6 (Heterochromatin / ZNF/ Repeats), 1 to E1 (Quiescent / No mark), and -1 to E5 (Quiescent / Weak repression). This allow to further increased the selection power as the top 1% features by variance ranking. We then used these top 1% variable features for subsequent analysis.

#### Dimension reduction and functional ontology analysis

We performed dimension reduction and visualization taking directly the categorical format of the above described selected features using multiple correspondence analysis (MCA). To explore the biological significance of the regions that contributed to the dimension that distinguishes the non-basal and basal clusters, the chromosome segments’ loading estimates to the Dim 2 were extracted from the MCA outputs and regions with a p-value < 0.05 for the loading estimate were included (n = 12,198). Genes mapped to Dim 2 contributing regions were pre-ranked by loading estimates for gene-set enrichment analysis (GSEA) by which we identified multiple biological gene sets / ontologies associated with Dim 2. The gene sets collections were retrieved from the Broad Institute Molecular Signature Database, spanning the H (hall mark gene sets), C2 (curated gene sets, eg. pathways), C3 (regulatory target gene sets), C5 (ontology gene sets, eg. Gene Ontology), C6 (oncogenic signature gene sets), and C8 (cell type signature gene sets) categories, using the msigdbr R package (27, 60).

As complementary exploration, we in the meantime performed dimensional reduction to the numeric transformed data of the selected features, using MDS. Similar to what was observed in MCA, the Dim 2 represents the dimension that distinguishes the basal versus non-basal samples, and Dim 1 separates cell lines from tumours, suggesting potential batch effect and/or *in vitro* culture-specific effect. We then adjusted for these latent effects to obtain a refined clustering (basically on Dim 2), using the MNN algorithm implemented in the fastMNN function of the batchelor R Bioconductor package (61).

We then calculated for the chromosomal segments (ie. features of the consensus sample-by-states matrix), the difference in the numeric chromatin state scores between basal and non-basal groups, named chromatin state score difference basal vs non-basal. A negative score difference indicates stronger activation in the non-basal group, and a positive one indicates stronger activation in the basal group. For subsequent function analysis, we performed expression quantitative trait locus (eQTL) mapping to refine the segments to the ones significantly linked with associated gene expression, and limited the analysis to the significant eQTL pairs (p-value < 0.05, n = 4,377). We then analysed the distribution of the chromatin state score difference of the segments corresponding to the luminal and basal cell type signature genes.

### Cell treatments, cell viability assay

For siRNA treatments, cells were reverse transfected using Lipofectamine RNAi max (Invitrogen) using 10 ng of siRNA (siRNA table in additional information).

For CRispR mutant cell lines production, RT112 and SD48 cells were plated at 80% confluence and the day after transfected with vectors expressing Cas9 an gRNA (VectorBuilder, see additional information) using Fugene HD transfection reagent. 48h post transfection, cells were selected using Puromycin (2µg/µL) during 4 days. After 2 weeks, clonal selection was performed using clonal dilution. FOXA1 mutation was assessed by Western Blot (anti-FOXA1 Abcam ab23738), PCR and genomic DNA sequencing.

Cell Viability was assessed in 96 well plates using CellTiter-Glo® Luminescent Cell Viability Assay (Promega).

### RNA extraction and sequencing

#### RNA extraction

Cell lines RNA were extracted using Qiagen RNeasy kit coupled with DNAse treatment. Tumours RNA were extracted using Triple extraction protocol.

#### Tumours RNA sequencing

RNA sequencing libraries: Kit Nugen. The pool of libraries was quantified using a qPCR method (KAPA library quantification kit, Roche). The sequencing was carried out using paired-end mode (PE100) on a Illumina Novaseq 6000 instrument, using a custom primer (provided into the Nugen kit) to initiate the Read 1 sequencing. The target number of reads was about 50 million paired-reads per sample.

#### Cell lines RNA sequencing

RNA sequencing libraries were prepared from 1µg of total RNA using the Illumina TruSeq Stranded mRNA Library preparation kit (Illumina) which allows to perform a strand specific RNA sequencing. A first step of polyA selection using magnetic beads is done to focus sequencing on polyadenylated transcripts. After fragmentation, cDNA synthesis was performed and resulting fragments were used for dA-tailing and then ligated to the TruSeq indexed adapters. PCR amplification was finally achieved to create the final cDNA library (12 cycles). The resulting barcoded libraries were then equimolarly pooled and quantified using a qPCR method

(KAPA library quantification kit, Roche). The sequencing was carried out using paired-end mode (PE100) on a Illumina HiSeq2000 instrument. The sequencing configuration was set to reach an average of 100 million paired-reads per sample.

#### Cell lines 3’RNA-seq (Lexogen 3’Seq )

RNA sequencing libraries were prepared from 200ng of total RNA using the QuantSeq FWD 3’mRNA Seq LEXOGEN Standard (CliniSciences). Libraries were prepared according to the manufacturer’s recommendations. The first step enables the synthesis of double strand cDNA, by revers transcription, using oligo dT priming. A qPCR optimization step was performed in order to estimate the most appropriate number of PCR cycles for library amplification. The resulting amplified and barcoded libraries were then equimolarly pooled and quantified using a qPCR method (KAPA library quantification kit, Roche). The sequencing was carried out using single-read mode (SR100) on an Illumina Novaseq 6000 instrument. The sequencing configuration was set to reach an average of 10 million reads per sample.

### RNA-seq analysis

RNAseq were aligned on genome hg19 using STAR with default parameters. Our RNA-seq as well as RNA-seq from public data repository integrated using Deseq2 default parameters and VST normalisation. 3’RNA-seq were analysed with Deseq2 and RPM normalisation.

### Assignment of MIBC and NMIBC subtypes

Gene expression data of the most tumour cases was previously generated and published (9, 22). We assigned consensus classes using the previously generated gene expression data using ConsensusMIBC (v1.1.0) R package (3) (Table S1). Given potential intra-tumour molecular heterogeneity, we also to verify the subtype in our ChIP-seq sampled tumour area using RNAseq from the same powder using the same ConsensusMIBC (v1.1.0) R package.

NMIBC samples (n=2) were classified using classifyNMIBC R Package (7).

### Regulons

The regulatory network was reverse engineered by ARACNe-AP (41) from human urothelial cancer tissue datasets profiled by RNA-seq from TCGA. ARACNe was run with 100 bootstrap iterations using all probe-clusters mapping to a set of 1,740 transcription factors. Parameters used were standard parameters, with Mutual Information p-value threshold of 10^-8^.

The VIPER (Virtual Inference of Protein-activity by Enriched Regulon analysis) (42) (R package viper 1.24), using the regulatory network obtained from ARACNE on urothelial cancer, and we computed the enrichment of each regulon on the gene expression signature using different implementations of the analytic Rank-based Enrichment Analysis algorithm.

### SE Correlation Network

To build SE driven correlation network, we first selected genes regulated by SE defined as TF in Lambert *et al.* (40). Next using TCGA regulon VIPER score, we calculated the mean regulon score by subtype (Luminal, Ba/Sq or Stroma-Rich and kept TFs with mean regulon > 2 or < -2 (n=75). We further restricted the list to TFs with a minimum expression correlation of 0.5 in TCGA to build correlation network using igraph.

### General bioinformatics, statistical analyses and public data

Plots and statistical analyses were performed in R software version 3.6.1, using ggpubr package. Wilcoxon and Kruskal-Wallis tests were used to test the association between continuous and categorical variables, for 2 categories or >2 categories, respectively. P-values <0.05 were considered statistically significant. Pairwise correlation of gene expression was calculated using Pearson coefficient and plotted using complexHeatmap R package. All gene expression heatmaps show mean-centered log2-transformed normalized counts of each represented gene.

TCGA-BLCA MIBC RNA-seq data were downloaded from TCGA data portal using TCGAbiolinks package (R), raw counts were normalized to account for different library size and the variance was stabilized with VST function in the DESeq2 R-package (62). TCGA-BLCA samples (n=404) were classified using the consensus system using consensusMIBC R package.

CCLE urinary tract cell line gene expression were downloaded from the DepMap portal (https://depmap.org/portal/download/). For consensus classification of CCLE bladder cancer cell lines, we adapted this classification considering only genes expressed by both tumours and cell lines. Ten cell lines were classified as Ba/Sq, all other were grouped as non-Ba/Sq.

MGHU3 RNA-seq bulk data were download from GEO (accession number, GSE171129).

#### Survival analysis

For Kaplan Meier survival analyses testing the association of gene expression and overall survival, we used http://tumorsurvival.org/index.html tool and divided the samples based on mean +/- sd. Log-rank P values were calculated to test the association between overall survival and low vs high expression groups.

#### Public scRNA-seq and Basal/Luminal signature

We downloaded the log2 TPM normalized gene expression of single cells from a Ba/Sq subtype MIBC tumour from the GEO database (accession number, GSM4307111). Initial quality control excluded genes expressed in less than 3 cells and cells with less than 200 genes. The top 2000 variable genes were used as features for subsequent PCA and the first 9 principal components were used for cell clustering and visualization by uniform manifold approximation and projection (UMAP) embedding. The marker genes of the luminal and basal tumour cells were calculated with Wilcoxon test based approach. The single cell RNA-seq data analyses were performed using the Seurat v4 package with default parameters unless otherwise specified.

Given the single-cell derived luminal and basal tumour cell signature was based on single-cell sequencing of primary *in vivo* tumour sample, and the *FOXA1* knock-out perturbation signature is likely limited to the genes regulated by *FOXA1* in an *in vitro* setting, it is important to adopt the cell subtype signatures to refine to the marker genes regulated by *FOXA1*, as a *FOXA1*-depdent luminal-basal plasticity signature which could be then used for further analyses involving *in vitro* transcriptomes. We first compared the perturbation and single-cell signatures by GSEA (perturbation DEG effect for ranking, and luminal / basal signatures as gene sets of interest) and found that in RT112 cell line, there was both significant enrichment of luminal signature in genes down-regulated in *FOXA1 KO* clones and significant enrichment of basal signature in genes up-regulated in *FOXA1 KO* clones. We then took the leading edge genes as the adopted *FOXA1*- depdent plasticity signature. As validation, this adopted signature showed similar enrichment in RT112 *FOXA1* KD assays and SD48 *FOXA1* KO assays, while the original cell type signature failed.

## Data Availability

The datasets supporting the conclusions of this article are available in the GEO repository under accession numbers:

GSEXXXXX for Tumours ChIP-seq

GSEXXXXX for Normal and Cancer cell culture ChIP-seq

GSEXXXXX for Tumours RNAseq

GSEXXXXX for Normal and Cancer cell culture RNA-seq

GSEXXXXX for functional assays 3’RNAseq

## Ethics approval and consent to participate

All patients provided written informed consent. All research in this study conformed to the principles of the Helsinki Declaration. All patients consent to participate

## Funding

The work was supported by grants from Ligue Nationale Contre le Cancer: (H.N-K., J.F., C.G., X-Y.M., L.C., F.D., C.K., Y.A., F.R., I. B-P.) as an associated team (Equipe labellisée), the "Carte d’Identité des Tumeurs" program initiated, developed and funded by Ligue Nationale Contre le Cancer, and a post-doctoral fellowship supporting H.N-K. J.F. was supported by the Fondation ARC pour la recherche sur le cancer, L.C. by FRM (Fondation Recherche Médicale) and X-Y.M. was supported by a fellowship from ITMO Cancer AVIESAN, within the framework of Cancer Plan. The work was also supported by a “PL-Bio” project funded by INCa (2016-146), the French Ministry of Education and Research, the CNRS, and the Institut Curie. ChIP-sequencing was performed by the GenomEast platform, a member of the ‘France Genomique’ consortium (ANR-10-INSB-0009).

## Conflict of Interest

The authors declare that they have no competing interests

## Abbreviation

SE: Super-enhancer
TF: Transcription Factor
NMIBC: non-muscle-invasive bladder carcinoma
MIBC: muscle-invasive bladder carcinoma
BLCA: Bladder Cancer Carcinoma
Ba/Sq: basal/squamous
LumU: luminal unstable
LumNS: luminal non-specified
LumP: luminal papillary

## Acknowledgements

We thank the NGS platform of Institut Curie, directed by Sylvain Baulande

## Authors’ contributions

H. N-K., J. F, performed the ChIP-seq and RNA-seq experiments.

C.K. established the NHU primary cells cultures.

Y. N. and T. L. provided clinical insight for analysis of the CIT cohort.

J.F. under Y.A. supervision performed histo-pathological analysis and E.G. the macrodissection of the tumors.

A.R. under D.G. supervision optimized RNA sequencing of primary tumour samples.

H. N-K. performed the functional experiments, siRNA, CRispR, cellular assays.

H. N-K, X-Y.M., C.G., J.F, T.Y., L.C., F.D, E.C. carried out the bioinformatics analysis. In particular, H.N-K. analysed ChIP-seq data, performed ChromHMM integration, SE identification with J.F., differential analysis, ontology/GSEA analysis, 3’RNA-seq analyses, and established coregulatory network with the help of J.F., F.D. and D.J. C.G. designed R scripts and participated to several analyses (GSVA, SE differential analysis, classifications). L.C. performed the regulon analysis. X-Y. M. analysed ChromHMM output, publicly available scRNA-seq, derived Basal/Luminal signature, 3’RNA-seq, performed GSEA analysis, optimized SE annotation. T.Y. under I.D. supervision processed the ChIPseq data. E.C. processed RNA-seq data of cell lines and centralized the data for bioinformatics analyses.

H. N-K., J.F., X-Y.M., C.G., D.J., I. B-P., F.R. wrote the manuscript.

The study was conceived by H. N-K., J.F. and supervised by Y.A., I.B-P. and F.R.

All authors gave critical insights and approved the final version for publication.

## Notes

### Competing Interest Statement

The authors have declared no competing interest.

