## Supplemental_Info_and_Figures for "Epigenomic mapping identifies a super-enhancer repertoire that regulates cell identity in bladder cancers through distinct transcription factor networks"

Additional material and methods: 2
Cell proliferation and Soft agar assays 2
siRNA 2
CRispR vectors and guides 2
Tools, Web sites, Data Base 2
Supplemental figures legends: 4
Figure S1: Epigenetic landscape of BLCA. 4
Figure S2: Chromatin State Map of BLCA 4
Figure S3: Identification of the bladder super-enhancer repertoire and subtype specificities 4
Figure S4: Super-enhancers regulate a network of candidate master transcription factors for bladder cancer subgroups 4
Figure S5: FOXA1 binding profile 4
Figure S6: FOXA1 regulates inflammation and cellular identity 5
Figure S7: ZBED2, a novel Basa 5
Supplemental Tables titles 6
Table S1: 6
Table S2: 6
Table S3: 6
Table S4 6
Table S5: 6
Table S6: 6
Table S7: 6
Supplemental figures 6

### Additional material and methods:

#### Cell proliferation and Soft agar assays

For cell proliferation assays, cells were siRNA reverse transfected with Lipofectamine RNAi max in 6 well plate. Every 24h post transfection and during 4 days, cells were counted using Malassez. Cells were then plated in soft agar and fixed after 21 days.

#### siRNA

| Target gene | Brand | Reference |
|---|---|---|
| siFOXA1_1 | Ambion | s6688 |
| siFOXA1_2 | Ambion | s6689 |
| siControl_1 | Ambion | 4390843 |
| siControl_2 | Ambion | 4390846 |
| siZBED2_1 | Ambion | s35780 |
| siZBED2_2 | Ambion | s35781 |
| siFOXA1 pool | Dharmacon | L-010319-00 |
| siControl pool | Dharmacon | D-001810-10- 05 |

#### CRispR vectors and guides

| Vector | Brand | Reference | |
|---|---|---|---|
| Cas9 + Scafold gRNA | VectorBuilder | VB161201-1064vnm | GTTTTAGAGCTAGAAATAGCAAGTTAAAA TAAGGCTAGTCCGTTATCAACTTGAAAAA GTGGCACCGAGTCGGTGC |
| Cas9 + FOXA1 gRNA (1) | VectorBuilder | VB161130-1033fmt | GCACTGCAATACTCGCCTTA |
| Cas9 + FOXA1 gRNA (2) | VectorBuilder | VB161130-1031bdf | CATGTTGCCGCTCGTAGTCA |

#### Tools, Web sites, Data Base

| Name | package | version | URL |
|---|---|---|---|
| R | | 4.1.0 | https://www.r-project.org/ |
| Rstudio | | 1.4.1106 | https://www.rstudio.com/ |
| | Diffbind | 2.16.0 | |
| | DESeq2 | 1.32.0 | |
| | Viper | 1.24.0 | |
| | fgsea | 1.18.0 | |
| | clusterprofiler | 4.0.0 | |
| | ReactomePA | 1.36.0 | |
| | pheatmap | 1.0.12 | |
| | ComplexHeatmap | 2.8.0 | |
| | ChIPseeker | 1.28.3 | |
| | FactoMineR | 2.4 | |
| | consensusMIBC | 1.1.0 | |
| | igraph | 1.2.6 | |

| | | | |
|---|---|---|---|
| | EnhanceVolcano | 1.10.0 | |
| | ggpubr | 0.4.0 | |
| | GSVA | 1.40.1 | |
| | TCGAbiolinks | 2.20.0 | |
| | msigdbr | 7.4.1 | |
| | classifyNMIBC | | |
| | batchelor | | |
| Easeq | | | https://easeq.net/ |
| Homer | | v4.11 | http://homer.ucsd.edu/homer/ |
| Cistrome | | | http://dbtoolkit.cistrome.org/ |
| Washu epigenome browser | | legacy | http://epigenomegateway.wustl.edu/legacy/ |
| ChromHMM | | v1.23 | http://compbio.mit.edu/ChromHMM/ |
| GraphPad Prism | | v8.4.3 | |
| GREAT | | | http://great.stanford.edu/public/html/ |
| Bedtools | | | https://bedtools.readthedocs.io/en/latest/ |
| ARACNe-AP | | | https://github.com/califano-lab/ARACNe-AP |

### Supplemental figures legends:

**Figure S1: Epigenetic landscape of BLCA.**

A) HES of primary tumors prior macrodissection.

B) WISP analysis of primary tumors RNAseq.

C) Signature analysis of primary tumors.

D) Macs2 Peaks number per sample for each tested histone mark

**Figure S2: Chromatin State Map of BLCA**

A) Transition parameters from CHromHMM state output.

B) ChromHMM output, example of Fold enrichment over genome categories (Tumour T391)

C) Pie chart illustrating chromatin state of all regions (entire genome), informative regions and top 1% varying regions by signed-rank.

D) MCA of Chromatin states on 1% most varying genome segments.

**Figure S3: Identification of the bladder super-enhancer repertoire and subtype specificities**

A) Ranked enhancer plots from ROSE algorithm based on H3K27ac signal in 4 tumors from our cohort: Regions corresponding to the dots to the right of the curve's inflexion point are considered super-enhancers by the algorithm. In tumors with a gene amplification (red box), the number of identified SEs is lower. The symbols of the gene closest to the top 10 ranked regions are shown at the right of each plot.

B) PCA of tumors samples for each histone marks within consensus SE regions.

C) Fold Change plots for Differential SE between Ba/Sq and LumP vs NHU (including NMIBC), significance by pvalue <0.05.

D) TCGA expression Heatmap of differential SE assigned genes (FDR<0.05, Clustering using Complete method).

**Figure S4: Super-enhancers regulate a network of candidate master transcription factors for bladder cancer subgroups**

A, B) Top55 TFs correlation of expression in TCGA (A), in CCLE (B).

C) Single cell RNAseq analysis of one Bladder Cancer tumour with both Basal and Luminal Population (GSM4307111). Right panels, expression of key markers in each single cell cluster.

**Figure S5: FOXA1 binding profile**

A) Genomic localisation of FOXA1 peaks.

B) Cistrome ChIPseq enrichment analysis of 5637 and SD48 FOXA1 overlapping peaks.

C) Homer motif enrichment analysis of FOXA1 SD48 specific FOXA1 peaks versus 5637 specific peaks.

D) Homer motif enrichment analysis of FOXA1 5637 specific FOXA1 peaks versus SD48 specific peaks.

**Figure S6: FOXA1 regulates inflammation and cellular identity**

A) Proliferation assay of SD48, RT112 and L1207 cell lines after FOXA1 KD with a pool of siRNA.

B) Soft agar assay of RT112 and L1207 upon FOXA1 KD.

C) FOXA1 expression upon siRNA in RT112 and SCaBER cell lines.

D) Msig Hallmark GSEA analysis of genes differentially regulated in siFOXA1 *vs* siControl samples (RT112 and SCaBER).

E) Genome browser view of STAT2 and CD274 loci. H2K27ac and FOXA1 ChIPseq and RNAseq in SD48 and 5637 cell lines.

F) Projection of TCGA tumors, RT112 and SCaBER siFOXA1 samples on Luminal/Basal adopted signature.

G) Western blot of SD48 and RT112 CRispR clones. Controls (Ct scramble), Mutants (FOXA1 sgRNA-1 or -2).

H) PCA of CRrispR clones 3'RNA-seq (Dim 2 and 3).

I) GSEA analysis of RT112 FOXA1 mutant clones on Basal and Luminal signature.

J) Venn diagram comparing differentially expressed genes in RT112 and SD48 FOXA1 CRispR clones.

**Figure S7: ZBED2, a novel Basal-associated TF involved in inflammation dampening**

A) Overall survival compared to ZBED2 expression in TCGA cohort (Mean +/- sd).

B) Top ZBED2 target genes predicted by ARACNE from TCGA regulon.

C) Statistical analysis of FOXA1 expression in 15 Pancreatic cell lines (GSE141607).

D) Expression correlation of ZBED2 with Interferon Gamma response (Hallmark) in TCGA cohort.

E) Viability assay of RT112 and SCaBER upon siRNA treatments (n=2).

F) 3'seq analysis of RT112 and SCaBER after siRNA treatment against ZBED2 and FOXA1.

#### Supplemental Tables titles

**Table S1:** Annotation table of samples indicating Tumour cell content, consensus classification and statistics

**Table S2:** ROSE SE table for each sample

**Table S3:** Consensus SE and analysis

**Table S4:** Reactome pathway analysis of Ba/Sq vs Luminal SEs

**Table S5:** H3K27ac peaks differential analysis inside SE and motif enrichment analyses using Cistrome and Homer

**Table S6:** DEG in siFOXA1 vs Controls

**Table S7:** DEG in CRispR mutant FOXA1 vs Controls

#### Supplemental figures

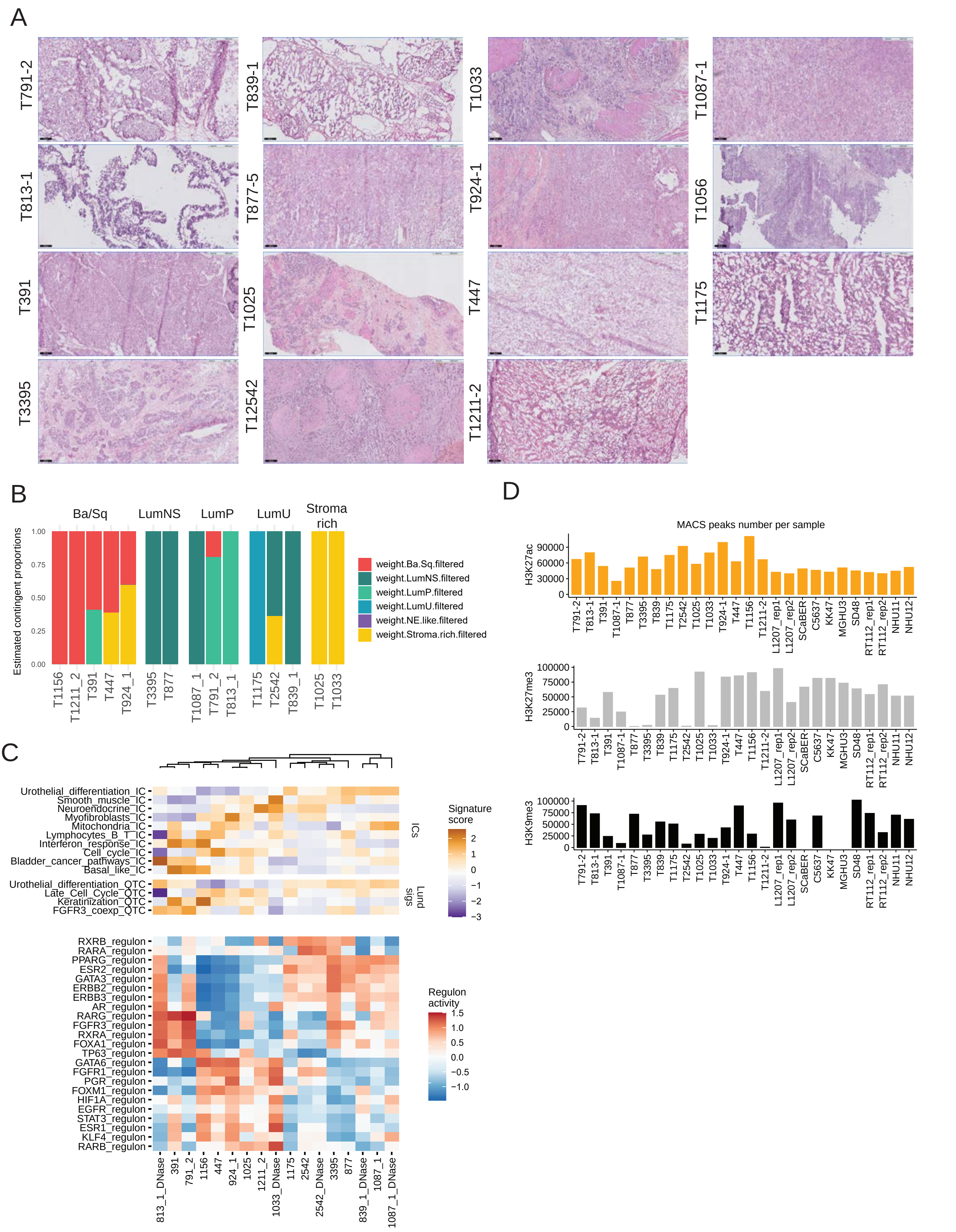


**Figure S1: Epigenetic landscape of BLCA.**
A) HES of primary tumor prior macrodissection. B) WISP analysis of primary tumors RNAseq.
C) Signature analysis of primary tumors. D) Macs2 Peaks number per sample for each tested histone marks

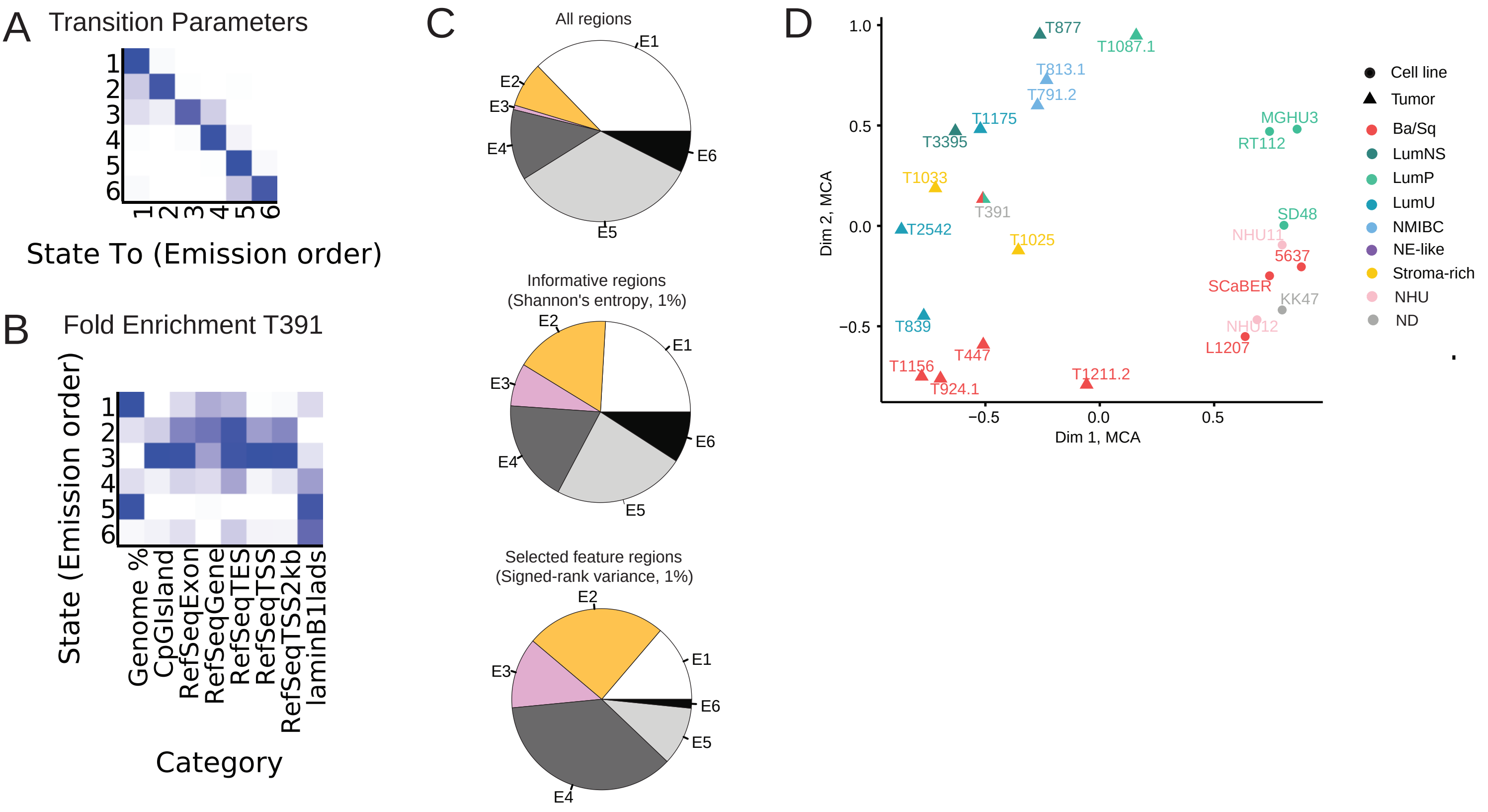


**Figure S2: Chromatin State Map of BLCA**

A) Transition parameters from CHromHMM state output.

B) ChromHMM output, example of Fold enrichment over genome categories (Tumour T391)

C) Pie chart illustrating chromatin state of all regions (entire genome), informative regions and top 1% varying regions by signed-rank.

D) MCA of Chromatin states on 1% most varying genome segments.

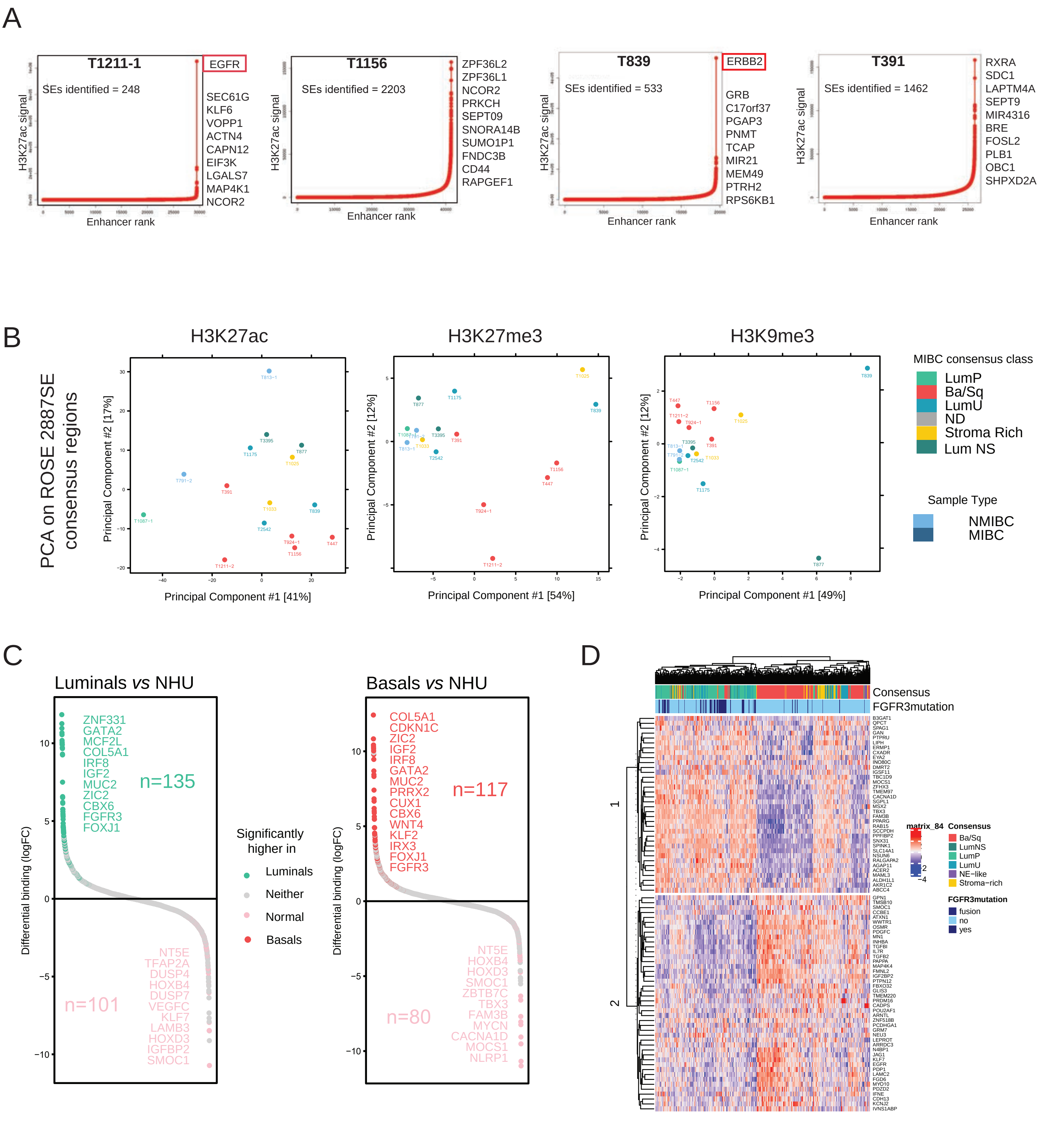


**Figure S3: Identification of the bladder super-enhancer repertoire and subtype specificities**
A) Ranked enhancer plots from ROSE algorithm based on H3K27ac signal in 4 tumors from our cohort: Regions corresponding to the dots to the right of the curve's inflexion point are considered super-enhancers by the algorithm. In tumors with a gene amplification (red box), the number of identified SEs is lower. The symbols of the gene closest to the top 10 ranked regions are shown at the right of each plot.
B) PCA of tumors samples for each histone marks within consensus SE regions. C) Fold Change plots for Differential SE between Ba/Sq and LumP vs NHU (including NMIBC), Significance by pvalue <0.05. D) TCGA expression Heatmap of differential SE assigned genes (FDR<0.05, Clustering using Complete method).

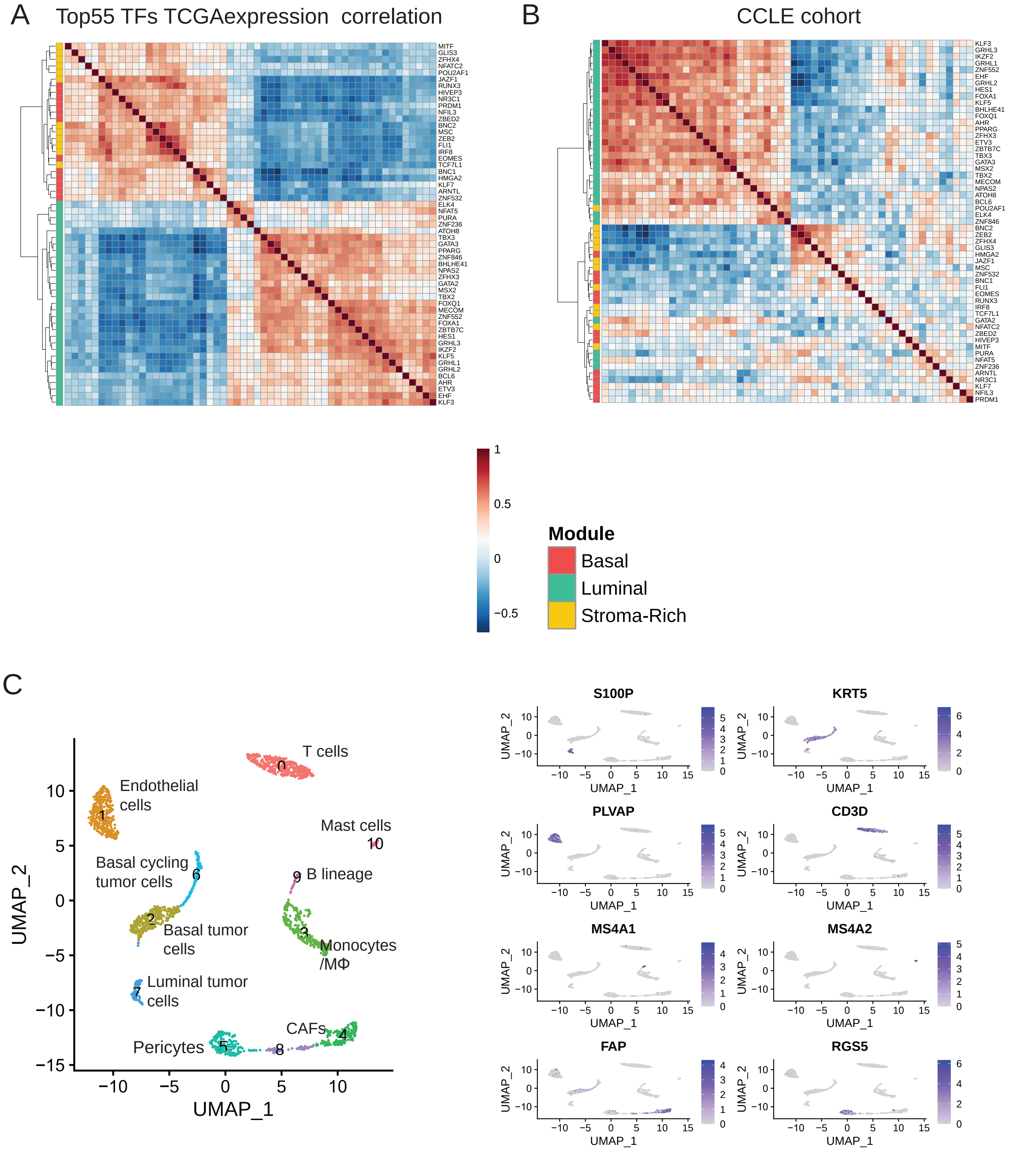


**Figure S4: Master Regulators**

A, B) Top55 TFs correlation of expression in TCGA (A), in CCLE (B).

C) Single cell RNA-seq analysis of one Bladder Cancer tumors with both Basal and Luminal Population (GSM4307111). Right pannels, expression of key markers in each single cell clusters.

A

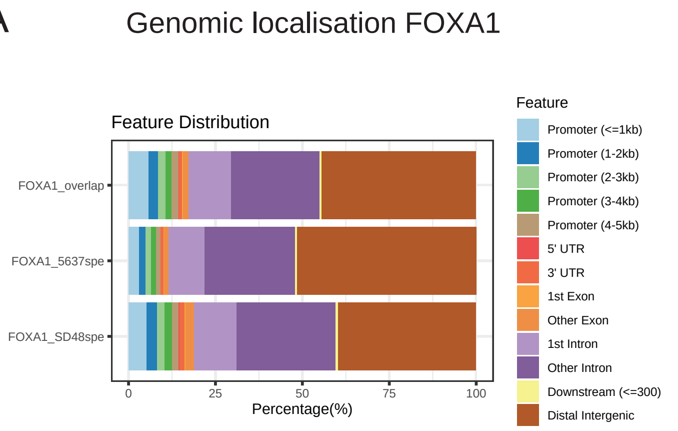


B

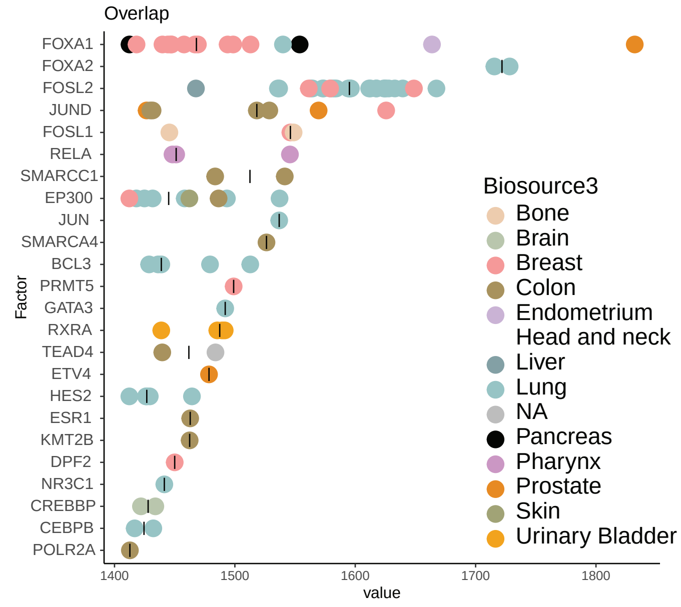


C

Homer motif enrichment FOXA1 SD48 specific peaks over 5637 specific peaks

| Motif Name | | Log P-value |
|---|---|---|
| Foxa2(Forkhead) | Liver-Foxa2-ChIP-Seq(GSE25694) | -3,38E+03 |
| GATA3(Zf) | iTreg-Gata3-ChIP-Seq(GSE20898) | -3,31E+03 |
| FOXA1(Forkhead) | LNCAP-FOXA1-ChIP-Seq(GSE27824) | -3,28E+03 |
| TRPS1(Zf) | MCF7-TRPS1-ChIP-Seq(GSE107013) | -3,12E+03 |
| FOXK1(Forkhead) | HEK293-FOXK1-ChIP-Seq(GSE51673) | -3,12E+03 |
| FOXA1(Forkhead) | MCF7-FOXA1-ChIP-Seq(GSE26831) | -3,06E+03 |
| FoxL2(Forkhead) | Ovary-FoxL2-ChIP-Seq(GSE60858) | -3,05E+03 |
| FOXK2(Forkhead) | U2OS-FOXK2-ChIP-Seq(E-MTAB-2204) | -2,93E+03 |
| Gata6(Zf) | HUG1N-GATA6-ChIP-Seq(GSE51936) | -2,78E+03 |

D

Homer motif enrichment FOXA1 5637 specific peaks over SD48 specific peaks

| Motif Name | | Log P-value |
|---|---|---|
| Fra1(bZIP) | BT549-Fra1-ChIP-Seq(GSE46166) | -1,27E+03 |
| Fos(bZIP) | TSC-Fos-ChIP-Seq(GSE110950) | -1,27E+03 |
| Fra2(bZIP) | Striatum-Fra2-ChIP-Seq(GSE43429) | -1,23E+03 |
| JunB(bZIP) | DendriticCells-Junb-ChIP-Seq(GSE36099) | -1,20E+03 |
| Atf3(bZIP) | GBM-ATF3-ChIP-Seq(GSE33912) | -1,20E+03 |
| BATF(bZIP) | Th17-BATF-ChIP-Seq(GSE39756) | -1,14E+03 |
| AP-1(bZIP) | ThioMac-PU.1-ChIP-Seq(GSE21512) | -1,11E+03 |
| Fosl2(bZIP) | 3T3L1-Fosl2-ChIP-Seq(GSE56872) | -1,06E+03 |
| Jun-AP1(bZIP) | K562-cJun-ChIP-Seq(GSE31477) | -9,24E+02 |

**Figure S5: FOXA1 binding profile**

A) Genomic localisation of FOXA1 peaks.

B) Cistrome ChIPseq enrichment analysis of 5637 and SD48 FOXA1 overlapping peaks

C) Homer motif enrichment analysis of FOXA1 SD48 specific FOXA1 peaks versus 5637 specific peaks.

D) Homer motif enrichment analysis of FOXA1 5637 specific FOXA1 peaks versus SD48 specific peaks.

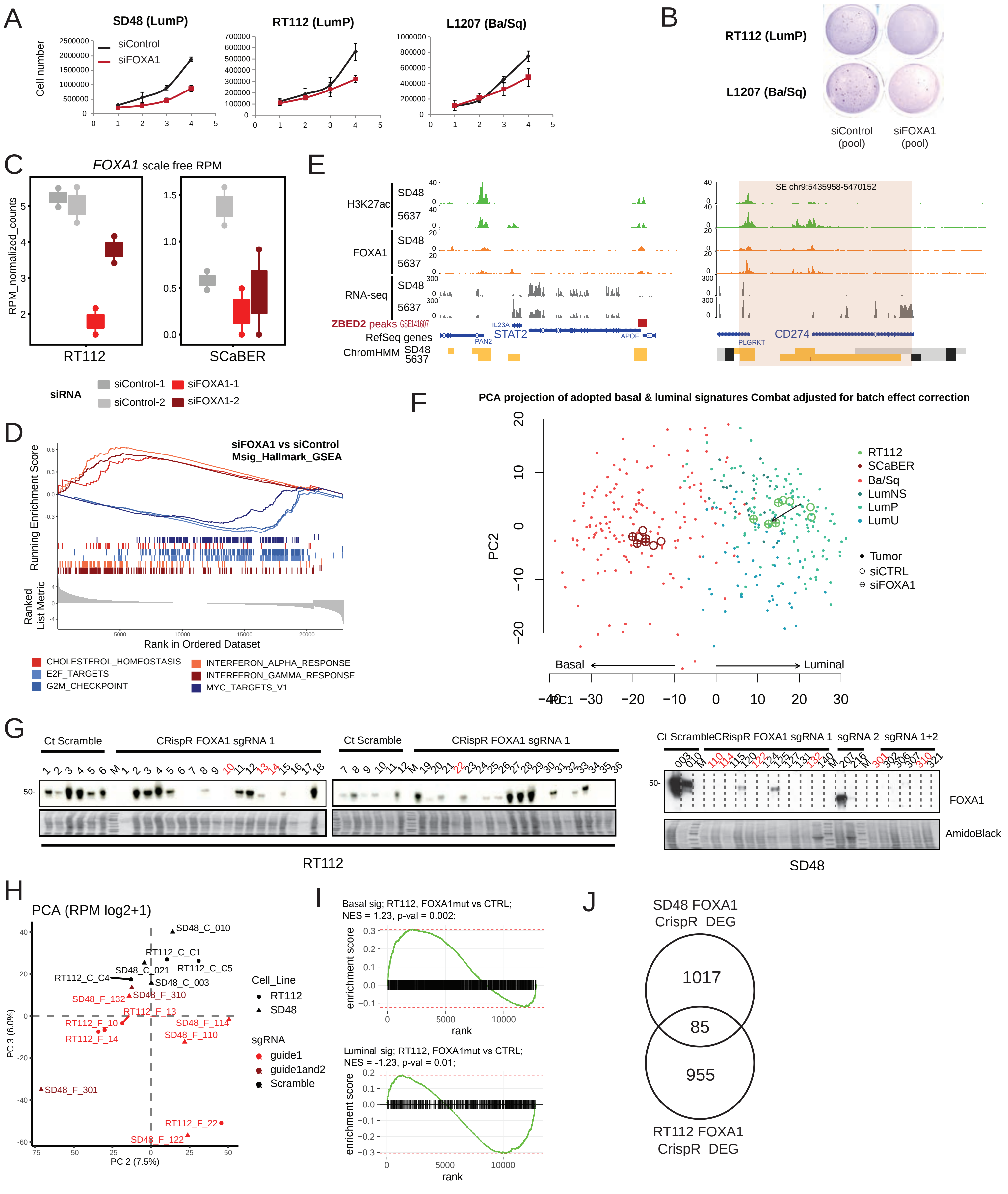

### Figure S6: FOXA1 regulates inflammation and cellular identity

A) Proliferation assay of SD48, RT112 and L1207 cell lines after FOXA1 KD with a pool of siRNA. B) Soft agar assay of RT112 and L1207 upon FOXA1 KD. C) FOXA1 expression upon siRNA in RT112 and SCaBER cell lines. D) Msig Hallmark GSEA analysis of genes differentially regulated in siFOXA1 *vs* siControl samples (RT112 and SCaBER). E) Genome browser view of STAT2 and CD274 loci. H2K27ac and FOXA1 ChIPseq and RNA-seq in SD48 and 5637 cell lines. F) Projection of TCGA tumors, RT112 and SCaBER siFOXA1 samples on Luminal/Basal adopted signature. G) Western blot of SD48 and RT112 CRispR clones. Controls (Ct scramble), Mutants (FOXA1 sgRNA 1 or 2). H) PCA of CrispR clones 3'RNA-seq(Dim 2 and 3). I) GSEA analysis of RT112 FOXA1 mutant clones on Basal and Luminal signature. J) Venn diagram comparing differentially expressed genes in RT112 and SD48 FOXA1 CRispR clones.

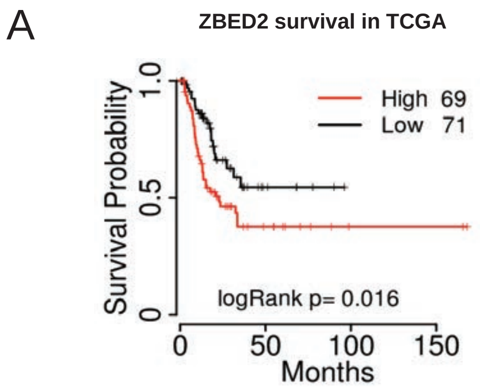


B

**Top ZBED2 target genes (Regulon ARACNE TCGA)**

| SYMBOL | mode | SYMBOL | mode |
|---|---|---|---|
| **FOXA1** | -0,998847 | PDCD1LG2 | 0,9999569 |
| **TBX3** | -0,997545 | **IL7R** | 0,9998151 |
| ZNF711 | -0,997167 | GBP5 | 0,9997805 |
| ZNF620 | -0,996492 | NCF2 | 0,999767 |
| GRM3 | -0,996456 | **CAV1** | 0,9997363 |
| ARFGEF3 | -0,996433 | CHST11 | 0,9997176 |
| ZNF10 | -0,996286 | MYO1G | 0,999701 |
| CYP4Z2P | -0,995916 | ADAM19 | 0,9996843 |
| ZFHX2 | -0,995471 | FCGR2A | 0,9996796 |

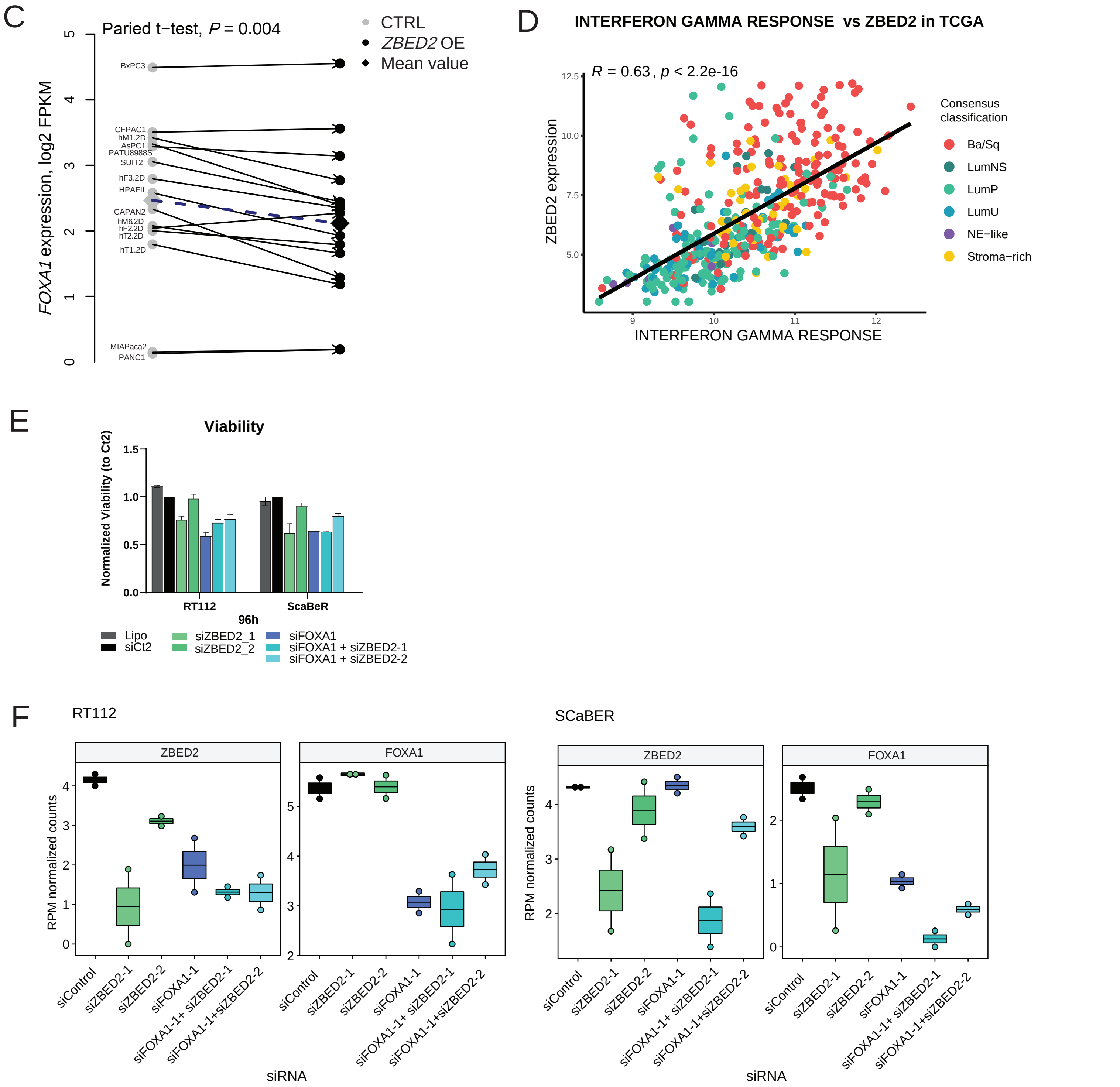


**Figure S7: ZBED2, a novel Basal-associated TF involved in inflammation dampening**

A) Overall survival compared to ZBED2 expression in TCGA cohort (Mean +/- sd). B) Top ZBED2 target genes predicted by ARACNE from TCGA regulon. C) Statistical analysis of FOXA1 expression in 15 Pancreatic cell lines (GSE141607). D) Expression correlation of ZBED2 with Interferon Gamma response (Hallmark) in TCGA cohort. E) Viability assay of RT112 and SCaBER upon siRNA treatments (n=2). F) 3'seq analysis of RT112 and SCaBER after siRNA treatment against ZBED2 and FOXA1.
